# Reduced Kv3.1 Activity in Dentate Gyrus Parvalbumin Cells Induces Vulnerability to Depression

**DOI:** 10.1101/784090

**Authors:** Lucian Medrihan, Gali Umschweif, Anjana Sinha, Shayna Reed, Katherina Gindinova, Subhash C. Sinha, Paul Greengard, Yotam Sagi

**Affiliations:** Laboratory for Molecular and Cellular Neuroscience, Rockefeller University, New York, N.Y, USA.

## Abstract

Parvalbumin (PV)-expressing interneurons are important for cognitive and emotional behaviors. These neurons express high level of p11, a protein associated with depression and action of antidepressants. Here we show that either specific deletion of p11 (p11 cKO) or chemogenetic inhibition in dentate gyrus (DG) PV neurons leads to anxiety-like behavior and susceptibility to depression in mice. DG PV neurons from p11 cKO mice showed reduced level and function of Kv3.1, and consequentially reduced capacity of high-frequency firing and altered short-term plasticity at synapses on granule cells. Activation of Kv3.1 or overexpression of the channel attenuated the vulnerability to depressive behavior.

## Introduction

Along with impairments in the glutamatergic signaling, recent imaging and biochemical studies suggest a reciprocal dysfunction of inhibitory signaling systems in the pathophysiology of stress related neuropsychiatric disorders. These include reduced levels of GABA in the blood, cerebrospinal fluid, and cortex of depressed patients ^1, 2^, as well as reduced levels of inhibitory neuronal markers including parvalbumin (PV), SNAP-25, and GABA receptor levels in the hippocampus from post mortem Major Depressive Disorder (MDD) and bipolar disorder subjects ^3, 4^. Preclinical studies support a link between stress and the dysfunction of the GABAergic signaling in the hippocampus. Early life stress in rats led to an increase and a decrease in hippocampal glutamate and GABA release, respectively ^5^, whereas α2 GABA-A receptors in distinct hippocampal microcircuits are required for the anxiety-reducing actions of anxiolytics ^6^. Among the inhibitory neuronal populations, the fast spiking parvalbumin (PV)-expressing cells play a central role in hippocampal function ^7^. These cells accommodate their firing frequency according to the excitatory input to modulate hippocampal granule and pyramidal cell output via both feedforward and feedback inhibitory modalities ^8^. Importantly, dysfunction of these cells is associated with neurological and psychiatric disorders, including schizophrenia ^9, 10^.

We recently identified an antidepressant role for p11, a member of the s100a calcium sensors ^11^. p11 in hippocampal cells is essential in mediating the response to antidepressants ^12–14^. In this brain region, p11 is highly enriched in PV and cholecystokinin (CCK) GABAergic interneurons ^12, 14, 15^. In CCK cells, p11 regulates the surface levels of the 5-HT1B serotonergic receptor to initiate the response to antidepressants, an effect mediated by disinhibition of PV cells ^16^. Still, the mechanism by which p11 directly regulates the function of PV cells remains unknown. In this study we identified a role for p11 in regulating PV hippocampal cell activity and in mediating the behavioral responses to novelty, stress, and antidepressant treatment, and elucidated the underlying mechanism. We show, in mice, that p11 in PV cells of the dentate gyrus (DG) regulates the level and activity of the Kv3.1 ion channel, by modulating the protein level and intracellular localization of the channel. Deletion of p11 from PV cells results in impairments in neuronal firing and short-term plasticity at PV to granule cell synapses, along with anxiety-like behavior, impaired recognition, susceptibility for depression and loss of the behavioral response to antidepressant treatment. Conversely, up regulation of Kv3.1β in DG PV cells, or activation of the channel using agonist induced anxiolytic behavior and resilience in response to chronic stress. Our data propose a central role for DG PV cells in mediating stress resilience and antidepressant response by dynamically accommodating inhibitory transmission in this brain circuit.

## Results

### Deletion of p11 from PV neurons induces susceptibility to depression and reduces their firing frequency

To examine if p11 in parvalbumin (PV)-expressing neurons plays a role in regulating emotional behaviors, mice with conditional deletion of p11 from PV cells (p11 cKO) were tested. In the open field test (OF), p11 cKO spent 20 ± 1.7% less time in the center of the arena relative to WT mice, an indication of anxiety-like behavior (Fig. 1a). In the elevated plus maze, another test for anxiety-like behavior, cKO mice spent 83 ± 6.3% less time exploring the open arm relative to WT mice (Suppl. Fig. 1). To test the possibility that p11 in PV cells mediates the adaptation in emotional behavior in response to novel stress, mice were subjected to subthreshold social defeat stress (SSDS, Fig. 1b). In the subsequent social interaction test (SI), all p11 cKO mice manifested social avoidance (Fig. 1c). This resulted in a 67 ± 9.0% reduction in the time spent interacting with an unfamiliar aggressor mouse, relative to that by WT mice (Fig 1d). Interestingly, p11 cKO mice did not manifest basal anhedonic or helplessness-like behaviors, as detected by the sucrose preference, the novelty suppressed feeding, the forced swim, and the tail suspension tests (Suppl. Fig. 1). However, the behavioral response to chronic antidepressant treatment as well as the ability to discriminate between novel and familiar objects were impaired in p11 cKO mice (Fig. 1e, Suppl. Fig. 1).

**Figure 1.**
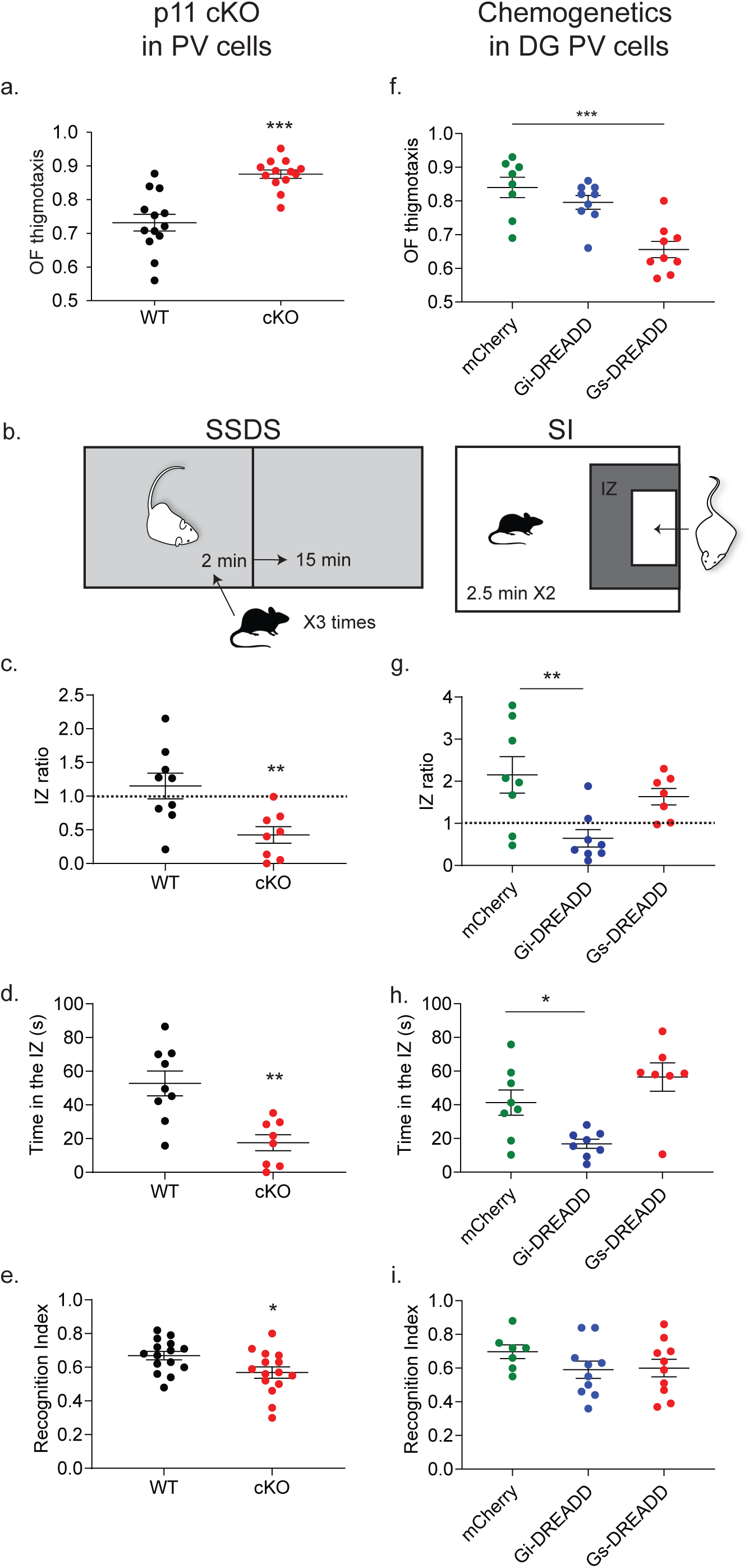
Deletion of p11 from PV neurons results in increased susceptibility to depression. **a**. Thigmotaxis behavior in the open field test (OF) in WT (black dots, n= 13 mice) and p11 cKO (red dots, n= 13). \*\*\**p*< 0.001 by unpaired *t*-test. **b**. Subthreshold social defeat stress (SSDS) included 3 defeat sessions. The social interaction test (SI) was conducted the next day. IZ, interaction zone. **c**. Ratio between the times spent in the IZ in the presence and absence of an unfamiliar aggressor in WT (n= 6) and p11 cKO mice (n= 6). \*\**p*= 0.007 by unpaired *t*-test. Dashed line represents social avoidance threshold. **d**. Time spent in the IZ in the presence of an unfamiliar aggressor. \*\**p*= 0.001 by unpaired *t*-test. **e.** Novel object recognition test (NOR), in WT (n= 15) and p11 cKO mice (n= 15). \**p*= 0.024 by unpaired *t*-test. **f.** OF thigmotaxis 3 weeks after DIO-DREADD virus injection to the DG in mCherry (n= 8), Gi-DREADD (n= 9), and Gs-DREADD (n= 9) mice. One Way ANOVA. *F* (2, 23) = 15.07, *P*<0.0001. \*\*\**p*<0.0001 by post hoc Bonferroni. **g.** Ratios of time spent in the IZ in mCherry (n= 8), Gi-DREADD (n= 8), and Gs-DREADD (n= 7) treated mice. One Way ANOVA. *F* (2, 20) = 6.43, *P*=0.007. \*\**p*= 0.004 by post hoc Bonferroni. **h.** Time spent in the IZ. One Way ANOVA. *F* (2, 20) = 9.21, *P*=0.002. \**p*=0.028 by post hoc Bonferroni. **i.** NOR in mCherry (n= 7), Gi-DREADD (n= 10), and Gs-DREADD (n= 10) mice. One Way ANOVA. *F* (2, 24) = 1.23, *P*= 0.319.

To test if an alteration in the activity of dentate gyrus (DG) PV cells might result in similar behavioral deficits, we utilized a chemogenetic approach and transfected Gi-, or Gs-DREADD into DG PV cells (Suppl. Fig. 2). In the OF, a single injection of clozapine n-oxide (CNO) in Gs-DREADD injected mice increased the time in the center of the arena by 22 ± 2.9% relative to that in mCherry controls (Fig. 1f). In the SSDS, acute CNO treatment induced social avoidance in 75% of the Gi-DREADD treated mice, along with a 59 ± 6.5% reduction in the time they interacted with an unfamiliar mouse, relative to that by mCherry-treated mice (Fig. 1g, h). In contrast, recognition of novel objects and helplessness behavior were not altered by chemogenetic manipulation of DG PV cells (Fig 1i, Suppl. Fig. 2). Taken together, these results suggest that p11 mediates resilience to depression by enabling the increase in the activity of DG PV cells in response to novel psychological stressors.

To investigate the impact of p11 deletion on the physiology of PV neurons, we used patch-clamp recordings in acute slices from the DG of WT and p11 cKO mice. The firing frequency of PV neurons from p11 cKO in response to injected current steps (100 pA) was lower than that in WT neurons (Fig. 2a, b), with no changes observed between genotypes in either membrane potential value or action potential properties (Suppl. Fig. 3). Kv3.1, a potassium channel highly specific and abundant in PV neurons has been shown to modulate the firing frequency of these cells ^17^. The Kv specific current was markedly reduced in p11 cKO mice in comparison with WT mice (Fig. 2c, d). In contrast, the amplitude of the HCN channel, another channel associated with the modulation of cell activity by p11 was not different in cKO vs WT (Suppl. Fig. 4). Interestingly, although the density of the Kv current at 50 mV was reduced by 43 ± 5.4 % in p11 cKO mice (Fig. 2e), the conductance of the channel was not different between genotypes (Fig. 2f), suggesting that p11 regulates the membrane expression of these channels without interfering with their functional properties.

**Figure 2.**
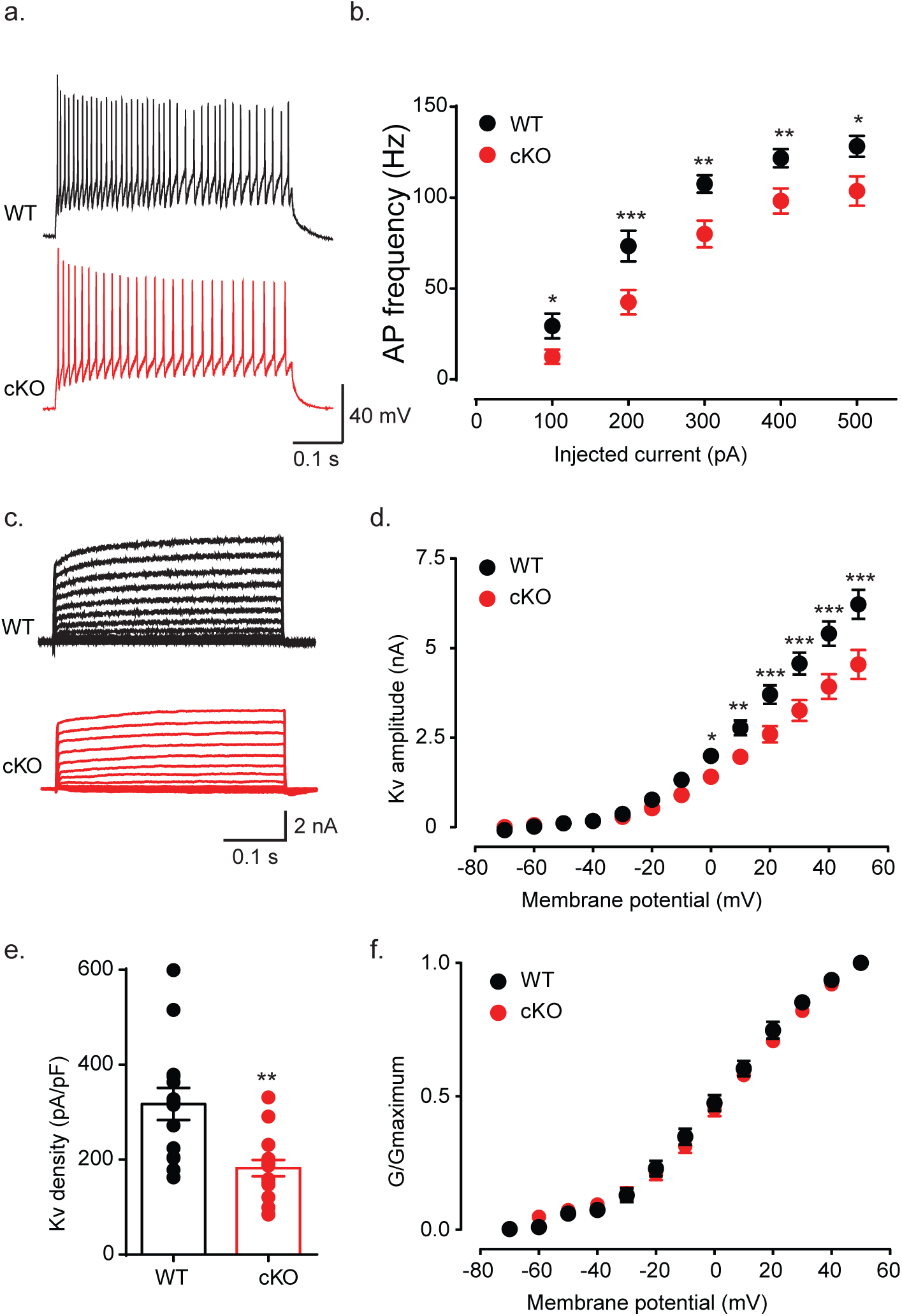
Deletion of p11 from PV neurons results in decreased action potential frequency and reduced Kv current amplitude. **a**. Representative traces of the firing response to a 200 pA step injection of current in PV neurons from WT and cKO mice. **b**. The firing frequency of PV neurons from WT (n = 16 neurons/ 10 mice, written as 16/ 10) and cKO (17/ 11) mice in response to 100 pA current steps injections. Two Way ANOVA. Genotype *F* (4, 134) = 73.43; P<0.0001, *p< 0.05, **p< 0.01, ***p< 0.001 by post hoc Uncorrected Fisher’s LSD test. **c.** Representative traces of Kv potassium currents evoked with 10 mV potential steps from −70 to +50 mV in WT and cKO mice. **d**. I-V curves showing the Kv amplitude in PV neurons from WT (16/ 8) and cKO (15/ 8) mice in response to increasing 10 mV potential steps. Two Way ANOVA. Genotype *F* (12, 377) = 183.7; P<0.0001, *p< 0.05, **p< 0.01, ***p< 0.001 by post hoc Uncorrected Fisher’s LSD test. **e**. Histograms showing the maximal density of Kv currents at +50 mV. \*\**p*=0.0011 by unpaired *t*-test. **f**. The relative conductance of Kv channels in PV neurons from WT (n = 13 / 7) and cKO (n= 12/ 7) mice in response to increasing 10 mV potential steps. Two Way ANOVA. Genotype *F* (1, 299) = 1.090, P=0.2972, post hoc Uncorrected Fisher’s LSD test.

### Deletion of p11 reduces Kv3.1 levels and abolishes the capacity of PV neurons to adapt to high-frequency firing

We next measured the protein level of Kv3.1 in hippocampal lysates from p11 WT and cKO mice. Western blot analysis confirmed a 41 ± 5.9% reduction in Kv3.1 protein level in p11 cKO mice, supporting the idea that p11 in PV cells modulates Kv3.1 function by regulating the expression level of the channel (Fig. 3a, b). To study whether p11 in PV cells regulates Kv3.1 synthesis, we measured the translated mRNA levels of the Kv3.1 α and β isoforms in hippocampal PV cells from WT and cKO mice, using translating ribosome affinity purification (TRAP). Semi-quantitative real time PCR (qPCR) analysis confirmed that Kv3.1β (the protein product of the *kcnc1a* transcript) is 3.3 fold more abundant than Kv3.1α (the product of *kcnc1b*, Fig. 3c). Importantly, no difference was detected in the ribosome-bound levels of either transcript in hippocampal PV cells between WT and cKO mice, supporting the idea that p11 regulates the channel by inhibiting the degradation of the protein (Fig. 3c).

**Figure 3.**
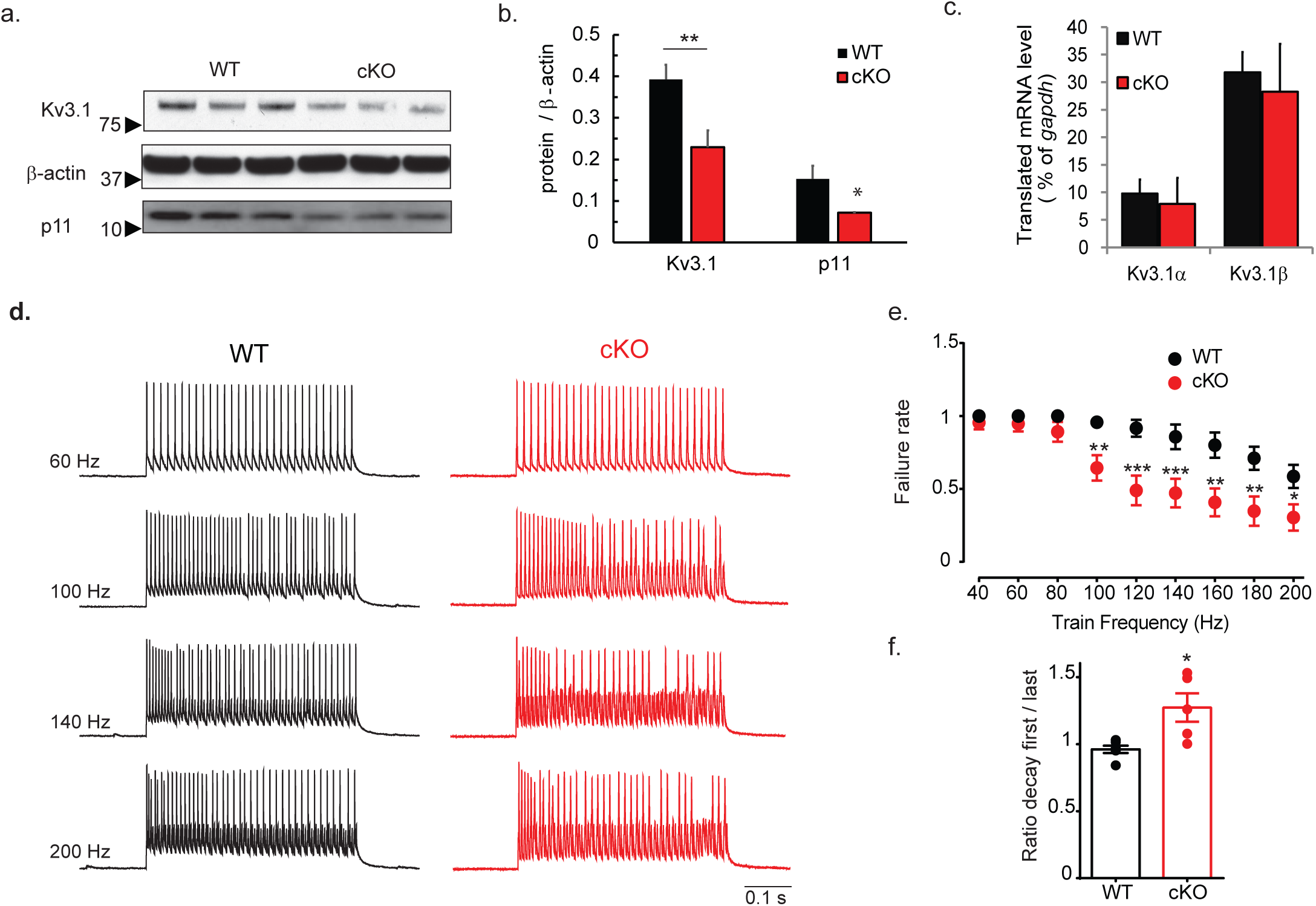
p11 regulates hippocampal Kv3.1 levels and mediates adaptation to high-frequency firing in DG PV neurons. **a**. Representative immunoblot from hippocampal lysates of WT and cKO mice. Arrows and numbers represent protein weights in KDa. **b**. Densitometry of the data presented in (**a**) in n= 3 animals per group. \**p*= 0.027, \*\**p*=0.006 by unpaired *t*-test. **c**. mRNA expression levels of Kv3.1 isoforms in hippocampal PV cells from WT (n= 7 replicates) and cKO (n= 4) PV*^TRAP^* mice. Unpaired *t*-test. **d**. Representative traces of action potentials evoked by trains of injected currents (1-2 nA, 1 ms) at different frequencies in PV neurons from WT and cKO mice. **e**. Failure rate (measured as ratio) induced in PV neurons from WT (n = 7/ 4) and cKO (n= 6/ 4) mice. Two Way ANOVA. Genotype *F* (10, 121) = 20.28; P<0.0001, \**p*< 0.05, \*\**p*< 0.01, \*\*\**p*<0.001 by post hoc Bonferroni. **f**. The ratio of the half-amplitude width between the last and the first action potential in the 100 Hz train in PV neurons from WT and cKO mice, \**p*=0.013 by unpaired *t*-test.

The most important physiological property of PV neurons is their ability to respond with high-frequency firing to inputs and this function is dependent on Kv3 channels in these neurons ^17^. We next measured the capacity of PV cells in WT and cKO mice to respond with action potentials induced by stimuli at increased frequencies (1 nA; 1ms; 10 to 200 Hz). We noticed that PV neurons from cKO mice have an increased failure rate above 100 Hz stimulations when compared with PV neurons from WT mice (Fig. 3d, e). The increased failure rate can be explained by a 32 ± 11.1 % increase in the width of the action potentials in a train in PV cKO (Fig. 3f), which reflects the reduced function of Kv3.1 in these neurons ^17, 18^.

### Presynaptic reduction of Kv3.1 disrupts short-term plasticity at the PV-granule cells synapse

Anxiety-like behavior and vulnerability to stress are likely to reflect an impairment in the network of neurons rather than cellular impairment of a single neuronal type. We measured the inhibitory and excitatory synaptic input on PV neurons from WT and cKO mice but we did not notice any difference between genotypes (Suppl. Fig. 5). Several studies have shown that Kv channels are highly expressed in the axon terminal of PV neurons where they inhibit neurotransmitter release and the GABAergic output to the principal cells ^19–21^. Therefore, we infected PV neurons in the DG with the excitatory opsin ChETA, and recorded their monosynaptic output on granule cells (GC) using optical stimulation and whole-cell patch-clamp (Suppl. Fig. 6). The amplitude of the postsynaptic GABAergic response in GC evoked by light stimulation of PV neurons increased by 119 ± 35.3% in the p11 cKO mice compared to WT mice (Fig. 4a, b). Bath addition of TEA (1mM), which blocks Kv channels, lead to a 67 ± 18.8% increase in the amplitude of the evoked postsynaptic current in PV-GC synapses from WT mice, but not in synapses from p11 cKO (Fig 4c, d), suggesting that presynaptic Kv channels are not functional in p11 cKO mice. Paired-pulse experiments showed a reduction in the ratio between the second and the first response in p11 cKO mice at short-time intervals (Fig. 4e, f). The block of Kv channels with 1 mM TEA decreased the paired-pulse ratio of WT mice to p11 cKO levels but had no effect on the paired-pulse ratio in the p11 cKO mice (Fig. 4g). Furthermore, a train of stimuli at 40 Hz, that depletes the ready-releasable pool at the PV-GC synapses showed a drastic effect in p11 cKO mice with more than 80% of the GABA vesicles being released after the first stimulation with respect to the ∼ 50% level noticed in WT synapses (Fig. 4h, i). The block of Kv channels with 1 mM TEA did not have any effect on the depletion kinetics of p11 cKO mice, but it brought the WT depletion kinetics to p11 cKO levels (Fig. 4i). Altogether, these data strongly suggest that presynaptic Kv channels are not functional in p11 cKO mice and, as a result, inhibition kinetics of GC by PV neurons are heavily altered.

**Figure 4.**
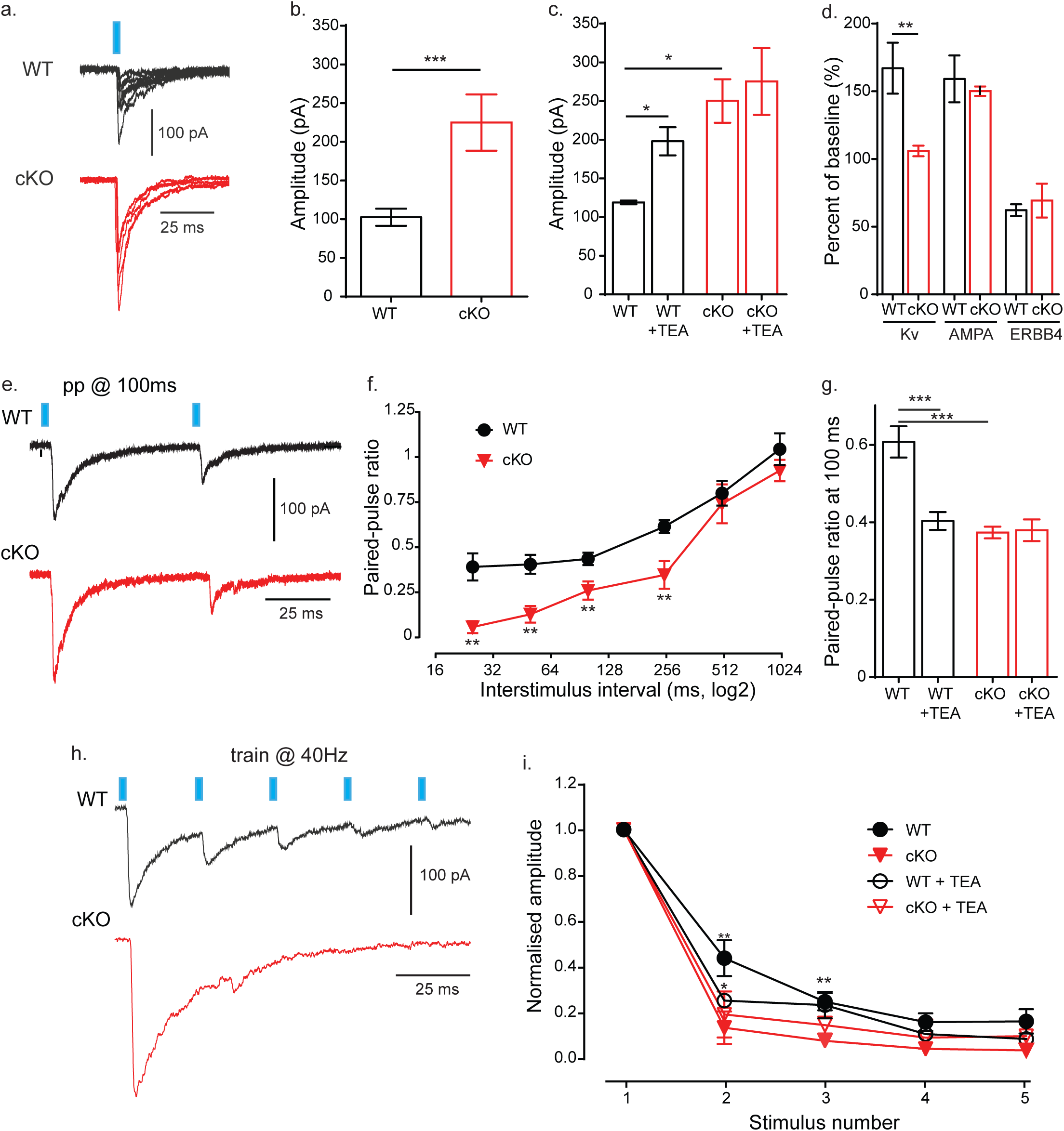
Reduction of Kv3.1 in DG PV neurons leads to loss of low-frequency synaptic filtering in PV-GC synapses. **a.** Representative monosynaptic GABAergic responses (**a**) evoked in GC neurons from WT and cKO mice by light stimulation of ChETA-AAV infected PV neurons. **b.** Mean amplitude in WT (n = 21 / 5) and cKO (n= 7/ 4), \*\*\**p*< 0.001 by unpaired *t*-test. **c**. Amplitude of monosynaptic GABAergic responses evoked by light in GC neurons from WT (n = 3/ 3) and cKO (n= 3, 3) mice before and after the bath application of 1 mM TEA. Two Way ANOVA. Genotype *F* (1, 4) = 8.123; P= 0.046, Treatment *F* (1, 4) = 16.88, P= 0.015, \**p*< 0.05 by post hoc Bonferroni. **d**. Changes in the monosynaptic GABAergic responses evoked by light in GC neurons from WT and cKO mice by different presynaptic modulators like Kv channels (blocked by 1mM TEA, n = 4/3 for WT and 3/3 for cKO), AMPA receptors (blocked by 10 mM CNQX, n = 4/3 for WT and 3/2 for cKO) and ERBB4 receptors (blocked by 10 µM PD158780, n = 4/2 for WT and 3/2 for cKO). One Way ANOVA. *F* (5, 14) = 14.76, P< 0.0001. \*\**p*= 0.0049 by post hoc Bonferroni. **e.** Representative traces of paired-pulse responses at 100 ms evoked in GC neurons from WT and cKO mice by light stimulation of ChETA-AAV infected PV neurons. **f**. Paired-pulse ratio of GABAergic responses evoked in GC neurons from WT and cKO mice by light stimulation at different interstimulation intervals in WT (n= 5-13/ 5) and cKO (n= 3-7/ 4). Two Way ANOVA. Genotype *F* (5, 58) = 42.80; P< 0.0001, \*\**p*< 0.01 by post hoc Bonferroni. **g.** Paired-pulse ratio at 100 ms evoked by light in GC neurons from WT (n = 3/ 3) and cKO (n= 3/ 3) mice before and after the bath application of 1 mM TEA. Two Way ANOVA. Genotype *F* (1, 4) = 11.27; P= 0.0284, Treatment *F* (1, 4) = 84.63, P= 0.0008, \*\*\**p*< 0.001 by post hoc Bonferroni. **h.** Representative traces of GABAergic responses evoked in GC neurons from WT and cKO mice by light stimulation at 40 hz of ChETA-AAV infected PV neurons. **i.** Normalized amplitude of GABAergic responses evoked in GC neurons from WT (n = 5/ 3) and cKO (n= 4/ 2) mice before and after the bath addition of 1 mM TEA. Two Way ANOVA. Genotype *F* (4, 55) = 203.3; P< 0.0001, Treatment *F* (3, 55) = 9.398, P< 0.0001, * *p*< 0.05, \*\*\**p*< 0.001 by post hoc Bonferroni.

### p11 regulates the protein level and intracellular localization of Kv3.1

Our studies indicate that p11 regulates the function of Kv3.1 in PV cells by regulating the cellular level of the protein. To study the detailed mechanisms by which p11 regulates Kv3.1, we measured changes in its level in N2A cells following transient co-transfection with either Kv3.1α or -β. Co-transfection with p11 increased the level of Kv3.1β and α proteins by 47 ± 13.0% and 431 ± 91.7% respectively (Fig. 5a-c), supporting the idea that p11 enhances the stability of the channel. Next, we stably transfected N2A cells with GFP-tagged Kv3.1β, and confirmed the molecular size of the tagged channel (Fig. 5d). Down regulation of p11 resulted in 31 ± 9.6% reduction in the protein level of Kv3.1β-GFP (Fig. 5e, f). Since p11 has been implicated in regulating the cell surface levels of several ion channels, it seemed likely that the regulation of Kv3.1 protein level by p11 is mediated via intracellular shuttling ^22–24, 25^. To visualize the changes in the intracellular localization of Kv3.1β by p11, we next applied immunocytochemistry and visualized the co-localization of GFP with established markers of the Golgi, endoplasmic reticulum (ER) and plasma membrane in fixed cells (Fig. 5g). Importantly, down regulation of p11 resulted in reduced localization of Kv3.1β to the Golgi apparatus by 11 ± 1.6%, while increasing its localization to the cell membrane by 47 ± 4.1% (Fig. 5h), supporting a role for endocytic shuttling by p11 in the regulation of Kv3.1.

**Figure 5.**
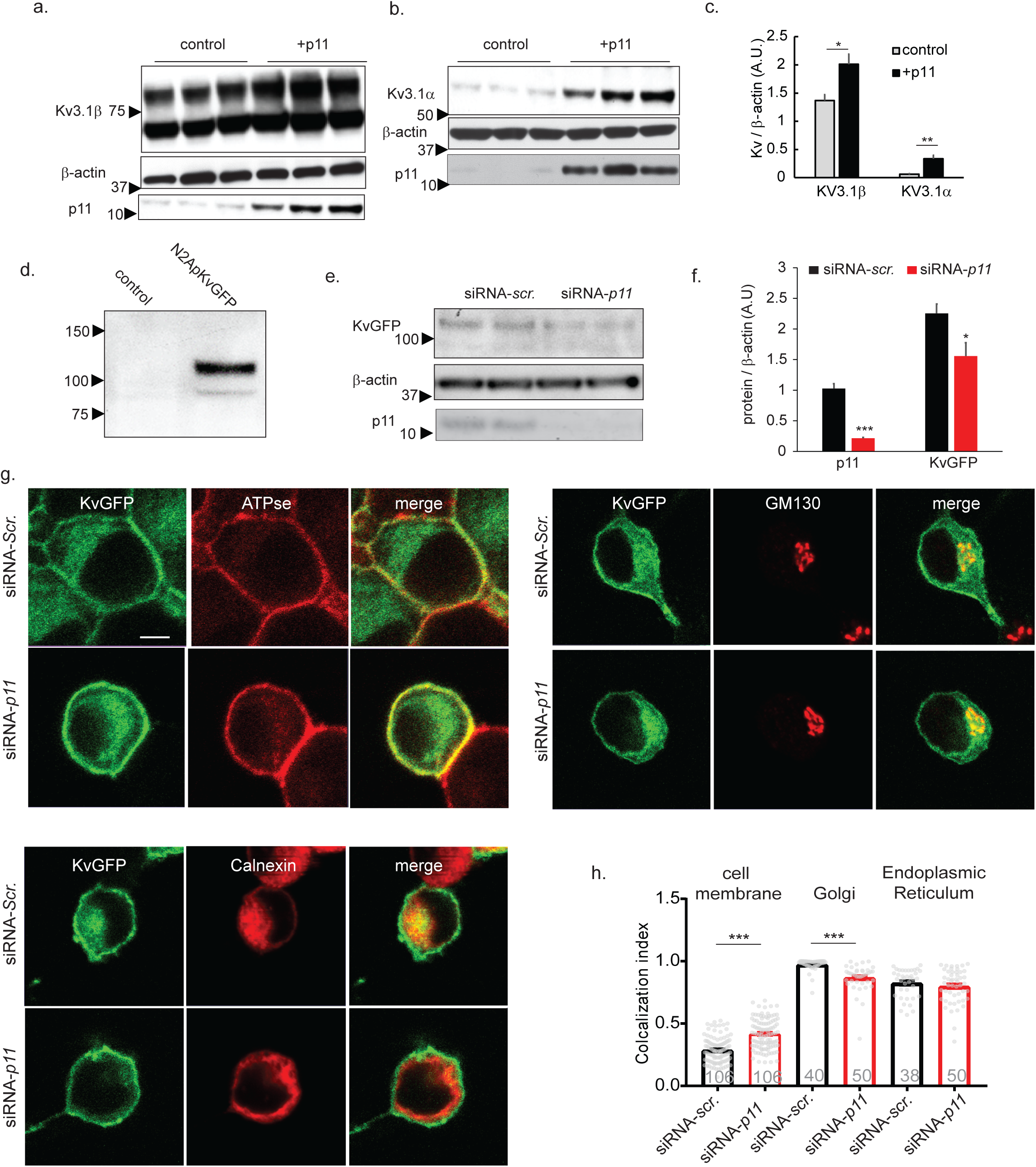
p11 regulates Kv3.1 shuttling. **a**. Representative immunoblot from N2A cell lysates after 48 hrs. of transient co-transfection of Kv3.1β with p11 or without (control). Arrows and numbers represent protein weights in KDa. **b**. Immunoblot from N2A lysates after 48 hrs. of co-transfection of KV3.1α with p11 or without (control). **c**. Densitometry of the protein levels in n=5 wells per group. \**p*= 0.031, \*\**p*= 0.001 by unpaired *t*-test. **d**. Immunoblot from representative lysates of N2a cell lines: Untransfected (control) or stably transfected with Kv3.1β-GFP. **e**. Cells were transfected with siRNA for p11 or control (siRNA-scr.) and lysed after 24 hrs. **f.** Densitometry of the protein levels in siRNA-p11 (n= 4 wells) and siRNA-scr (n= 4 wells). \**p*= 0.039, \*\*\**p*< 0.001 by unpaired *t*-test. **g**. Representative immunocytochemical images depicting co-localization between GFP and ATPase 1A1 (left), GM130 (right) and calnexin (bottom) in fixed Kv3.1β-GFP N2A cells transfected with either siRNA-scr. or siRNA-p11 for 24 hrs. Scale Bar, 5 μm. **h.** Co-localization ratio between GFP and the organelle markers. Dots represent individual cells and numbers inside the bars indicate the numbers of inspected cells. \*\*\**p*< 0.001 by unpaired *t*-test.

### Genetic upregulation or chemical activation of Kv3.1 induce resilience to chronic stress

To test if a direct upregulation of Kv3.1 in DG PV cells might induce anxiolytic response to novelty, we next generated AAV bearing a cre recombinase-dependent Kv3.1β, and injected it to the DG of PV-Cre mice (Fig. 6a). In the OF, mice overexpressing (O/E) Kv3.1β in DG PV cells spent 24 ± 3.4% more time in the center of the arena, relative to PV-Cre mice that were injected with the GFP control virus (Fig. 6b). Interestingly, O/E of Kv3.1β in p11 cKO mice did not improve thigmotaxis, further supporting the idea that in the absence of p11 Kv3.1β is not functional in DG PV neurons (Fig. 6b). We next subjected Kv3.1β O/E mice to 10 days of chronic social defeat stress (CSDS). None of the Kv3.1β-O/E mice manifested social avoidance in the SI test (Fig. 6c), and Kv3.1β-O/E mice spent 61 ± 21.5% more time interacting with an unfamiliar mouse, relative to that spent by the GFP-O/E mice.

**Figure 6.**
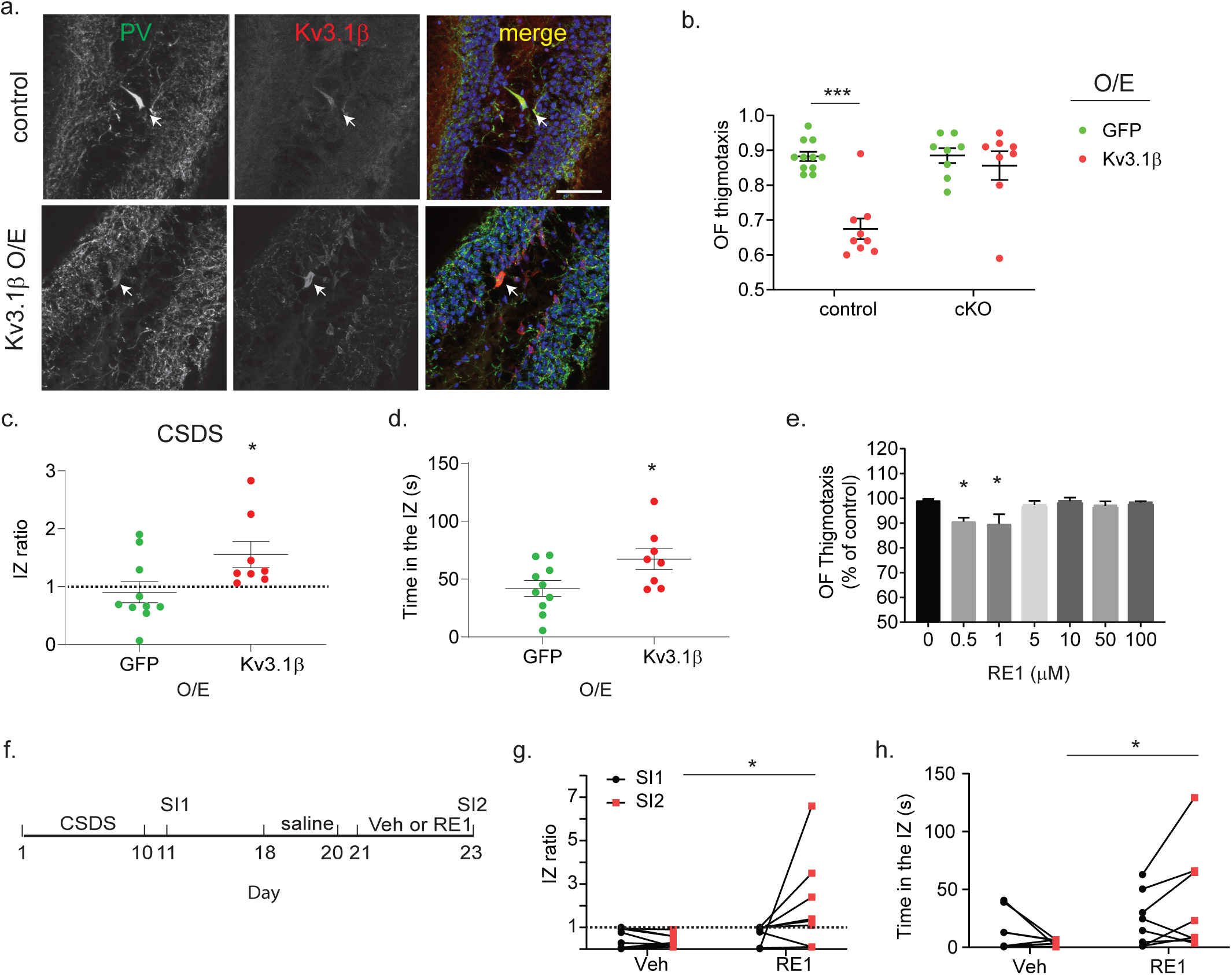
Activation of Kv3.1β induces resilience response. **a**. Representative immunohistochemical images showing Kv3.1β immunolabeling in DG PV cells from control mouse or in PV-Cre mouse after overexpression (O/E) of Kv3.1β. Scale bar, 50 μm. **b**. OF thigmotaxis after O/E of either GFP or Kv3.1β in DG PV cells, in PV-Cre (control, n= 11 GFP, and n= 9 Kv.3.1β) and p11 cKO (n= 8 GFP and n= 8 Kv.3.1β). Two Way ANOVA. *F* Genotype X AAV (1, 32) = 11.21; P= 0.0021, Genotype *F*= 11.78, P= 0.017; AAV *F*= 19.53, P= 0.001. \*\*\**p*< 0.001 by post hoc Bonferroni. **c**. Ratios of time spent in the IZ after CSDS in PV-Cre mice with O/E of GFP (n= 10) or Kv3.1β (n= 8) in DG PV cells. \**p=* 0.034 by unpaired *t*-test. **d**. Time spent in the IZ. \**p=*0.037 by unpaired *t*-test. **e**. WT mice were treated for 3 days with RE1 (0-100 μM in 100μl intraperitoneally) or with saline (control, n= 5 mice per group). OF thigmotaxis was determined 30 min after the third injection. One Way ANOVA. *F* (6, 28) = 4.1, P= 0.043. \**p*< 0.05 vs 0 μM by post hoc Bonferroni. **f-h**. Stress sensitive mice (IZ ratio ≤ 1) were identified in SI test (SI1) after CSDS, and were treated for 3 days with saline (100 μl intraperitoneally) and for additional 3 days either with RE1 (500 nM, 100 μl intraperitoneally) or Veh. The second SI test (SI2) was conducted 30 min after the last injection. **g**. Ratios of time spent in the IZ in n= 8 mice per group. \**p=* 0.049 by unpaired *t*-test. h. Time spent in the IZ in n= 8 mice per group. \**p*= 0.046 by unpaired *t*-test.

We then tested the anxiolytic and the antidepressant effects by RE1, an activator of the Kv3.1 channel, (Suppl. Fig. 7) ^26^. In the OF test, three days of treatment with 0.5 or 1 μM of RE1 resulted in 9 ± 1.5% and 10 ± 3.9% increase in the respective times spent in the center of the arena, relative to that by the vehicle-treated mice (Fig. 6e). In the SI test, three days of 0.5μM RE1 treatment resulted in a 450 ± 202.9% reduction in social avoidance in stress-sensitive mice, relative to that by the vehicle treated mice (Fig. 6f, g), along with a 867 ± 431.3% increase in the time they spent interacting with an unfamiliar mouse (Fig. 6h), supporting the idea that activation of Kv3.1 in DG PV cells may improve resilience to stress.

## Discussion

Here we describe a role for p11 in regulating the level and function of Kv3.1 channels in PV neurons of the hippocampus. GABAergic dysfunction has been proposed in depression, but a role for DG PV cell dysfunction in mediating the susceptibility for depressive behavior has not been previously shown. Using chemogenetics, we found that activation of DG PV cells induces an anxiolytic effect, whereas their inhibition increased the susceptibility to depressive behavior after subthreshold social stress. These results are in line with a recent study showing that inhibition of ventral DG GC exerts stress resilience in mice ^27^. Further, we found that Gi-DREADD mediated inhibition of DG PV cells did not induce effects in the tail suspension and object recognition tests. Similar to our results, Zou et al. reported that chemogenetic activation of DG PV using Gq-DREADD induced anxiolytic response but did not induce depressive-like behavior in TST or altered novel object recognition ^28^. Furthermore, we identified that the activation of Kv3.1 ion channels is a key mechanism to induce DG PV cell activity and resilience in response to stress. In light of these results it is expected that activation of DG PV neurons might be translated into a novel therapeutic approach in depressive states, as well as in other stress-related pathologies including anxiety and post-traumatic stress disorders.

### p11 regulates ion channels in different cell types via diverse mechanisms

In PV neurons, Kv3.1 is localized to both dendritic spines and axonal terminals, where it respectively mediates cell firing and neurotransmission ^17^. Notably, both of these functions were impaired in p11 cKO mice. Deletion of p11 resulted in reduced Kv3.1 protein levels in hippocampal lysates, and a similar effect was found in transfected cells following down regulation of p11, supporting the idea that the cellular level of the Kv3.1 protein is the target of p11. Previous studies identified a role for p11 in regulating the function of a variety of ion channels ^22–24 25^. Very recently we reported that p11 regulates the function of the HCN2 ion channels in cholinergic neurons from the nucleus accumbens ^29^. The effect by p11 on HCN2 function was mediated via transcriptional regulation of its gene expression. In line with this, it was previously suggested that the behavioral response to antidepressant treatment is dependent on p11 via the regulation of transcription in hippocampal cells, an effect that was attributed to the interaction between the protein complex p11/Annexin A2 and the chromatin remodeling factor SMARCA3 ^14^. Still, whether the p11/Annexin A2 /SMARCA3 complex plays a role in PV cell function after antidepressants remain unknown. Here we confirmed that p11 regulates the Kv3.1 protein level, and that this effect is not transcriptionally mediated. Furthermore, the expression level of the HCN2 channel was not altered in hippocampal PV cells in the absence of p11, as suggested by the unaltered function of the channel in the cKO mice. Taken together, these studies strongly support the idea that diverse mechanisms by which p11 regulates ion channel function, and suggest that the downstream channel targets of p11 are cell-type specific. Still, the downregulation of p11 in cultured cells increased the localization of Kv3.1 to the cell membrane but reduced it from the Golgi, supporting the idea that p11 regulates the internalization of the channel from the cell membrane. p11 is essential for the trafficking of diverse ion channels, including: TASK1, ASIC1a, NaV1.8, as well as that of metabotropic G-protein coupled receptors ^11, 16, 23, 25, 30, 31^. It was suggested that association of p11 with ion channels masks their endoplasmic reticulum retention signal ^22^. Together, our data supports the idea that impaired protein shuttling leads to reduced stability of the Kv3.1 ion channel in vivo, and that the role of p11 in the shuttling of the channel may involve endocytosis. Future studies should determine the exact mechanism by which p11 regulates the cellular localization of Kv3.1 and whether this effect is dependent on interaction with Annexin A2 or another binding partner of p11.

### Multiple roles for DG PV cells in mood-related behaviors

p11 is highly associated with Major Depressive Disorder and response to antidepressants ^11, 12^. In the hippocampus, p11 is enriched in PV and CCK cells as well as in mossy cells ^12, 14, 15, 32^. Here we show that p11 in PV cells is essential for mediating resilience in response to stress as well as regulating anxiolytic and cognitive responses to novelty. The behavioral deficits in p11 cKO were mimicked by chemogenetic inhibition of DG PV cells, suggesting that p11 activates PV neurons to mediate emotional as well as cognitive modalities of DG functions. The impairment in recognition in p11 cKO mice is in line with a previous report showing a similar dysfunction in mice bearing the constitutive deletion of p11 ^33^. Chemogenetic inhibition of DG PV cells mimicked the impairments in anxiolytic response in p11 cKO mice and the increased susceptibility to depression, whereas the recognition of familiar objects was unchanged by DREADD manipulation, suggesting that cognitive and emotional responses to novelty are regulated by DG PV cells in molecular mechanisms that are similar but in spatially different regions of the hippocampus. In line with this, bidirectional chemogenetic manipulation of the dorsal hippocampus mediated opposing effects on subthreshold novel object recognition^34^.

The deletion of p11 from PV cells did not induce anhedonia or helplessness behaviors. This is in line with previous reports showing that deficits in reward and motivation-related behaviors are found in mice with constitutive deletion of p11 or in those with conditional deletion in the cholinergic neurons of the nucleus accumbens ^11, 35, 36^. The idea that differences in mood-related behaviors are regulated by p11 in different cell types and brain regions is supported by the fact that the behavioral response to SSRIs is impaired in mice with deletion of p11 from hippocampal CCK, mossy cells and layer 5a cortical cells, but not cholinergic accumbal cells ^16, 32, 35, 37^. Taken together, the current study identifies a unique role for p11 in DG PV cells in mediating anxiolytic response and resilience for depression, as well as in regulating the behavioral response to chronic antidepressant treatment.

### Opposing regulation of Kv3.1 activity is required before and after chronic SSRIs

PV interneurons play a critical role in neuronal networks for complex processes, such as learning and memory, cognition, and emotional behavior, whereas disruption of their function has been associated with several mental illnesses and most consistently with epilepsy and schizophrenia^8, 9^. In PV neurons, Kv3 channels provide the rapid repolarization of action potentials during a very brief interspike interval, allowing high frequency firing rates ^17, 38^. Moreover, in terminals of presynaptic neurons, Kv channels contribute to neurotransmitter release evoked by a presynaptic action potential ^19–21^. We show here that alterations in both of these functions of Kv channels in PV neurons lead to anxiogenic response and susceptibility to depression. These findings are in line with recent data suggesting that the pathophysiological effects of PV neurons in epilepsy and schizophrenia may be related also to the dysfunction of Kv3.1. For example, a recurrent de novo mutation in Kv3.1 that suppresses the current amplitude when assembled into heteromers with WT Kv3.1 results in progressive myoclonus epilepsy, an inherited disorder that causes tonic-clonic seizures ^39^. In schizophrenia, the levels of Kv3.1β are significantly reduced in the prefrontal and parietal cortex of untreated schizophrenic patients and these levels were normalized by antipsychotic medication, suggesting that these agents restore Kv3.1β levels to those in controls ^40^. Moreover, it was recently shown that mice expressing truncated Disrupted-in-Schizophrenia 1 (Disc1), which mirrors a high-risk gene for psychiatric disorders including schizophrenia and depression show depressive-like behavior ^41, 42^. This behavioral deficit was correlated with reduced number of PV interneurons, disrupted synaptic input and output and abnormal limbic network oscillations in the low-gamma range, further supporting a major role for these neurons in pathological emotional behavior^41^.

Interestingly, in contrast to the activation of Kv3.1 that is required for resilience to depression, the action of antidepressant drugs may result in reduced activity of this channel. We recently found that the behavioral response to chronic SSRIs is dependent on the serotonergic 5-HT5A receptor signaling in DG PV cells, and that this signaling pathway mediates delayed inhibition of Kv3.1 channel function. This inhibitory effect is detected only after chronic SSRI treatment, and was mediated by an upregulation of the phosphorylation level of Ser-503 Kv3.1β ^43^. Indeed, our results strongly support the idea that the activity of Kv3.1 in DG PV cells is highly modulated during the course of chronic SSRI treatment, with maximal activity during the initiation of the treatment and subsequent reduction that requires delayed activation of the 5-HT5A receptor ^43^. Two recent findings support the idea that DG PV cell activity is dynamically changed during the course of the SSRI treatment. First, the initial SSRI treatment is associated with increased DG PV cell activity. This induction of activity was found to be mediated by 5-HT1B heteroreceptors on DG CCK cells, and their activation lead to reduced disinhibition of DG PV cells ^14^. Secondly, we recently showed that activation of DG PV cells after chronic SSRI treatment attenuated the behavioral response to SSRIs ^43^. Taken together, our studies suggest that changes in DG PV cell activity during chronic SSRI treatment correlates with the changes in the function of the Kv3.1 channel. The opposing regulation of Kv3.1 activity before and after chronic SSRIs, is initially mediated by p11, which is required for the initial demand in Kv activity, and later by the 5-HT5A receptor and its signaling which mediate the subsequent down regulation of the channel activity. Our current and previous findings suggest that the regulation of the ionic mechanisms that allow PV neurons to accommodate high-firing frequencies is essential for the DG microcircuit to cope with stress and novelty. Future studies should identify the details underlying the dynamic regulation of DG PV cell function in resilience to depressive behavior and after chronic antidepressant treatment.

## Acknowledgements

We are grateful to Jodi Gresack for valuable suggestions on behavioral studies, to Jerry Cheng for suggestions on imaging studies, to Elisabeth Griggs for the artwork, to Debra Poulter for proofreading, and to Jinah Lee, Zintis Inde, Chelsea Daniels and Katia George for technical assistance. This work was supported by The JPB foundation (P.G., and S.S.), The Leon Black Family Foundation (P.G.), and by the United States Army Medical Research Acquisition Activity Grant W81XWH-14-0390 (Y.S.).

## Author contributions

L.M., G.U., S.S., P.G. and Y.S. designed experiments and discussed results. L.M. performed and analyzed electrophysiology recordings, G.U designed, performed and analyzed social defeat studies. S.R. performed and analyzed biochemical and imaging studies. A.S synthetized RE1 with the assistance of K.G. Y.S designed and analyzed the molecular and behavioral studies. L.M and Y.S., wrote the manuscript.

## Legends to Supplementary Figures

**Supplementary Figure S1.**
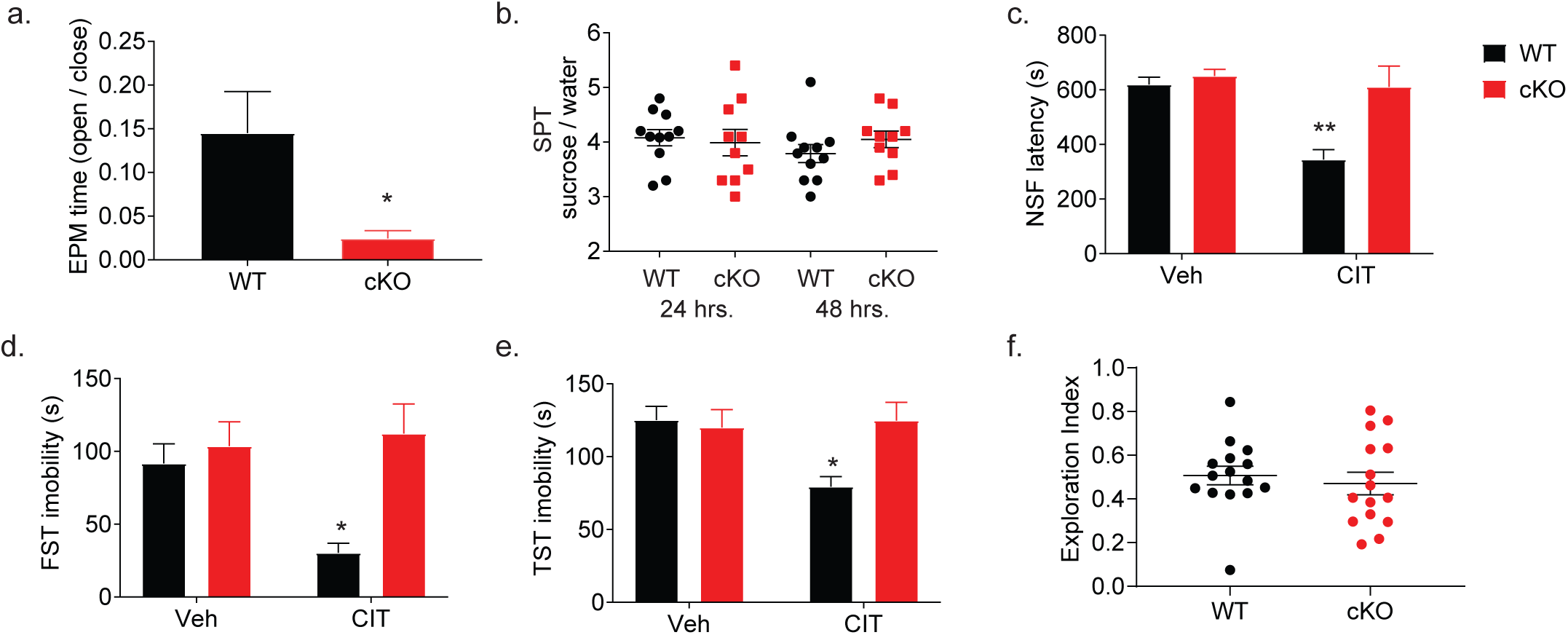
Depressive and anhedonic behaviors in p11 cKO. **a.** Ratio between the time spent in the open and closed arms in the elevated plus maze test (EPM) in WT (n= 9) and cKO (n= 9). \**p*= 0.024 by unpaired *t*-test. **b.** Sucrose preference test (SPT) in WT mice and cKO (n= 11, 10), after 24 and 48 hrs. of consumption. Unpaired *t*-tests. **c-e**. Effect of 14 days of citalopram (CIT, 10 mg/kg, intraperitoneally) or vehicle (Veh, 100 μl saline) on depressive-like behaviors. **c**. Latency to approach food pellet in the novelty suppressed feeding test (NSF) in WT mice (n= 7 Veh, 7 CIT) and cKO (n= 7 Veh, 8 CIT). Two Way ANOVA. *F* Genotype X treatment (1, 25) = 5.8, P= 0.024. \*\**p*=0.004 vs. WT Veh by post hoc Bonferroni. **d.** Immobility time in the forced swim test (FST) in WT (n= 7 Veh, 7 CIT) and cKO (n= 7 Veh, 7 CIT). Two Way ANOVA. *F* Genotype X treatment (1, 24) = 5.4, P= 0.030. \**p*= 0.040 vs. WT Veh by post hoc Bonferroni. **e**. Immobility time in the tail suspension test (TST) in WT (n= 8 Veh, 8 CIT) and cKO (n= 7 Veh, 8 CIT). Two Way ANOVA. *F* Genotype X treatment (1, 27) = 5.8, P= 0.023. \**p*=0.019 vs. WT Veh by post hoc Bonferroni. **f**. Exploration in the NOR in WT (n= 15) and p11 cKO (n= 15). *p*= 0.589 by unpaired *t*-test.

**Supplementary Figure S2.**
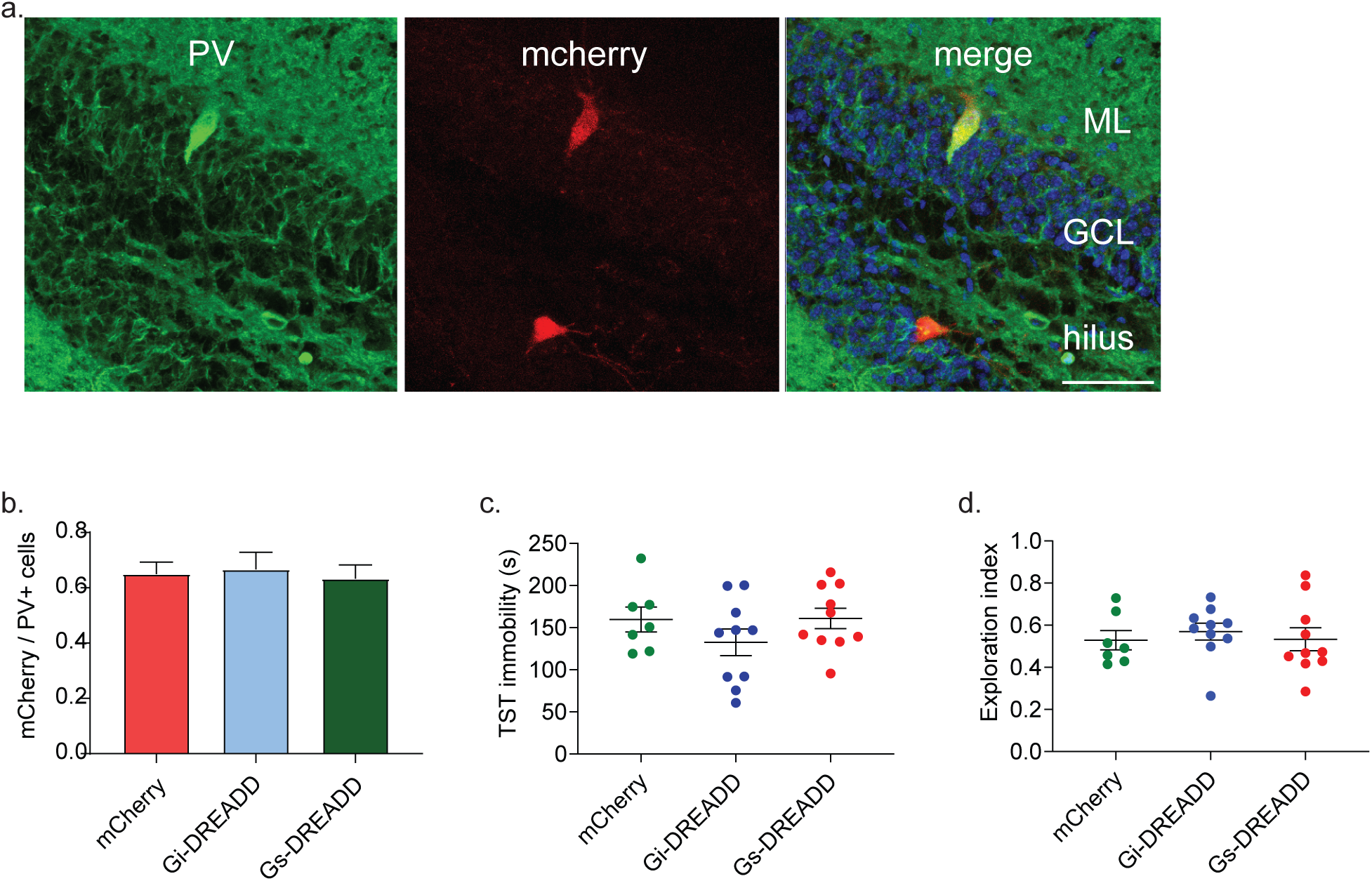
Chemogenetics manipulation in DG PV cells does not affect recognition and helplessness. **a.** Representative immunohistochemical images confirming co-expression of mCherry in DG PV cells after AAV injection of mCherry. Scale bar, 50 μm. ML, molecular layer; GCL, granule cell layer. **b**. Quantification of the fraction of PV cells infected with DREADD in mCherry (n= 6), Gi-DREADD (n= 6), Gs-DREADD (n= 6) mice. One Way ANOVA. *F* (2, 15) = 0.103, P= 0.902. **c**. Immobility time in the tail suspension test (TST) after chemogenetic manipulation in mCherry (n= 7), Gi-DREADD (n= 10), and Gs-DREADD (n= 10) mice. One Way ANOVA. *F* (2, 24) = 1.30, *P*= 0.29. **d.** Exploration in the NOR in mCherry (n= 7 mice), Gi-DREADD (n= 10 mice), Gs-DREADD (n= 10 mice). One Way ANOVA. *F* (2, 24) = 0.221, P= 0.803.

**Supplementary Figure S3.**
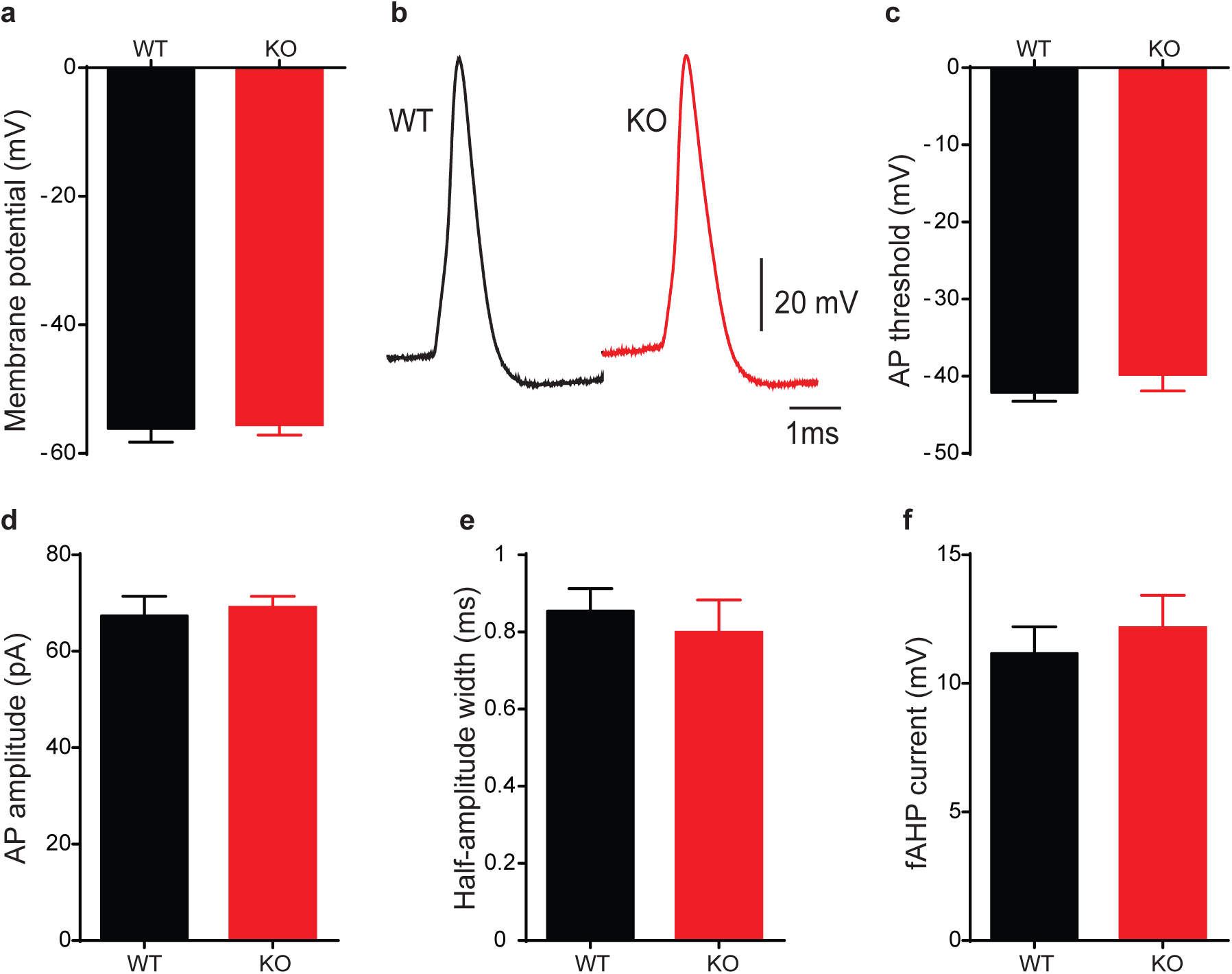
Basal physiological properties of PV neurons are not different between WT and cKO mice. **a.** Resting membrane potential is not significantly different between PV neurons from WT and cKO mice. **b**. Representative action potentials from WT and cKO DG PV neurons. **c-f**. Action potential properties such as AP threshold (**c**), AP amplitude (**d**), half-amplitude width (**e**) and the fast after hyperpolarization current (fAHP, **f**) show no significant difference between PV neurons from WT and cKO mice. n = 16 neurons/ 8 mice for WT and 15 neurons/ 8 mice for cKO. Unpaired *t*-tests.

**Supplementary Figure S4.**
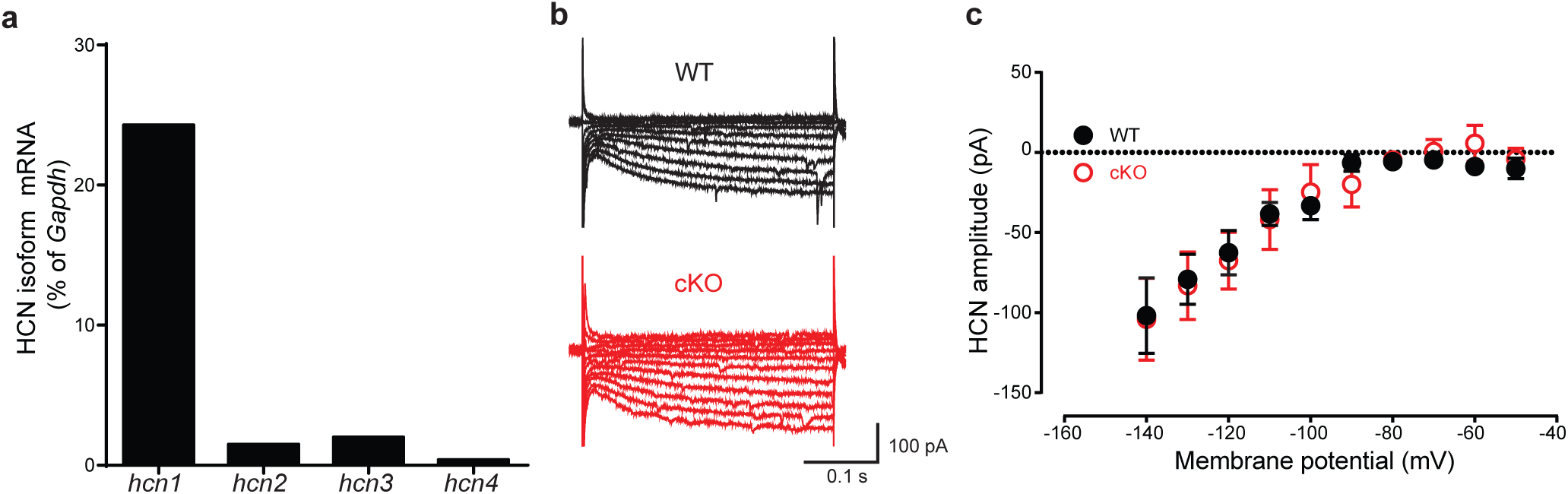
HCN currents in PV neurons are not different between WT and cKO mice. **a.** Bar graph summary of mRNA expression levels of the HCN isoform genes in PV hippocampal neurons from p11 WT PV*^TRAP^* mice. **b**, **c**. Representative traces of HCN currents (**b**) and the I-V curves (**c**) showing the HCN amplitude in PV neurons from WT and cKO mice in response to increasing 10 mV potential steps from −140 mV to −50 mV (n = 5/ 3 for WT and 3/ 2 for cKO). Unpaired *t*-tests.

**Supplementary Figure S5.**
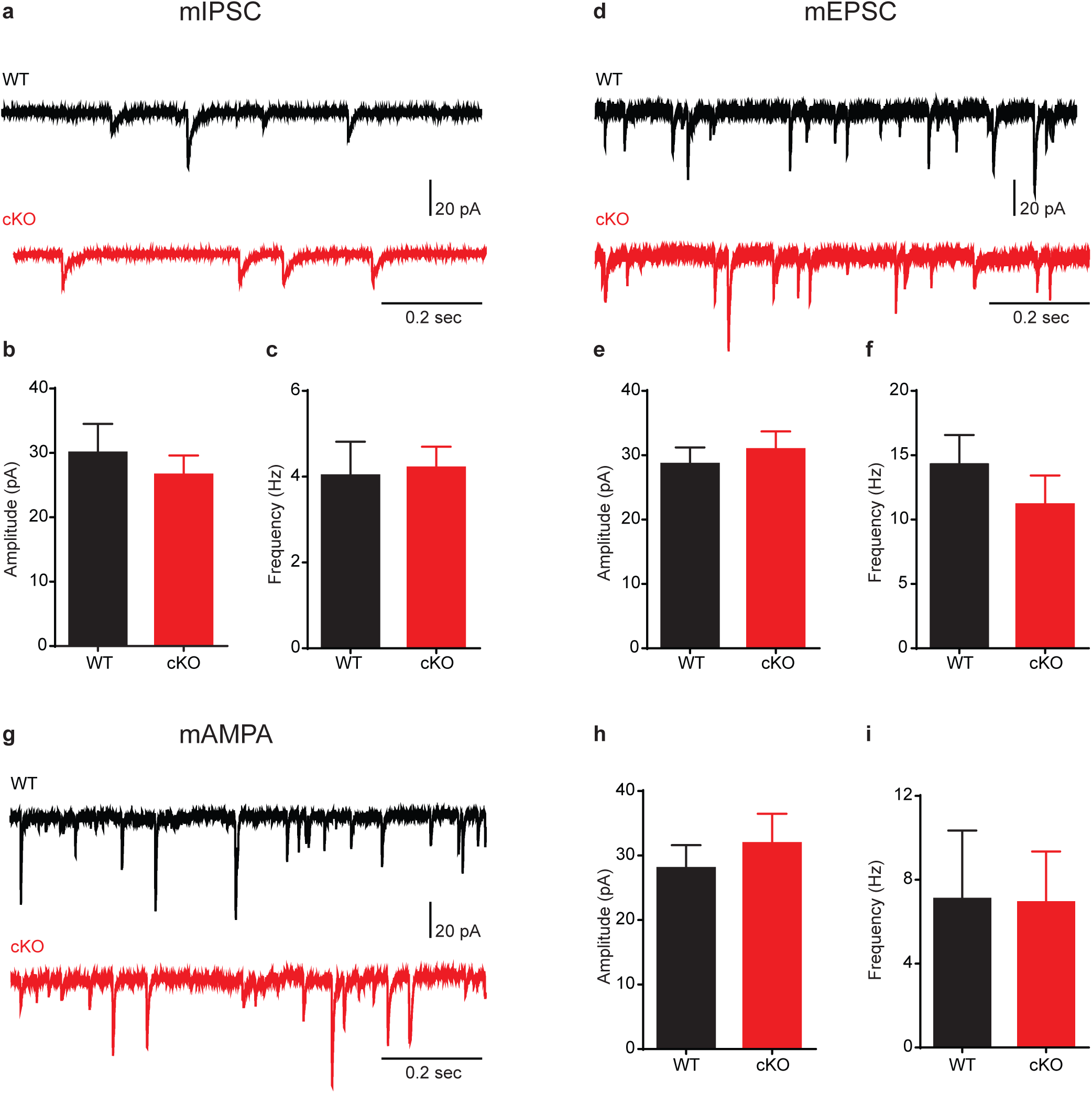
Inhibitory and excitatory synaptic input on DG PV neurons is not significantly different between WT and cKO mice. **a.** Representative traces of miniature inhibitory postsynaptic currents (mIPSCs) in PV neurons from WT (upper rows) and cKO mice (lower rows) measured in 1 µM TTX, 50 µM APV and 10 µM CNQX. b, c. Histograms showing that neither the amplitude (b) nor the frequency (c) of mIPSCs is changed between PV neurons from WT and cKO mice (n = 4/ 2 for WT and 6/ 3 for cKO). d. Representative traces of miniature excitatory postsynaptic currents (mEPSCs) in PV neurons from WT (upper rows) and cKO mice (lower rows) measured in 1 µM TTX and 30 µM bicuculline. e, f. Histograms showing that neither the amplitude (e) nor the frequency (f) of mEPSCs is changed between PV neurons from WT and cKO mice (n = 9/ 3 for WT and 14/ 4 for cKO). g. Representative traces of miniature excitatory postsynaptic AMPA currents (mAMPAs) in PV neurons from WT (upper rows) and cKO mice (lower rows) measured in 1 µM TTX, 50 µM APV and 30 µM bicuculline. h, i. Histograms showing that neither the amplitude (h) nor the frequency (i) of mAMPAs is changed between PV neurons from WT and cKO mice (n = 5/ 2 for WT and 5/ 2 for cKO). Unpaired *t*-tests.

**Supplementary Figure S6.**
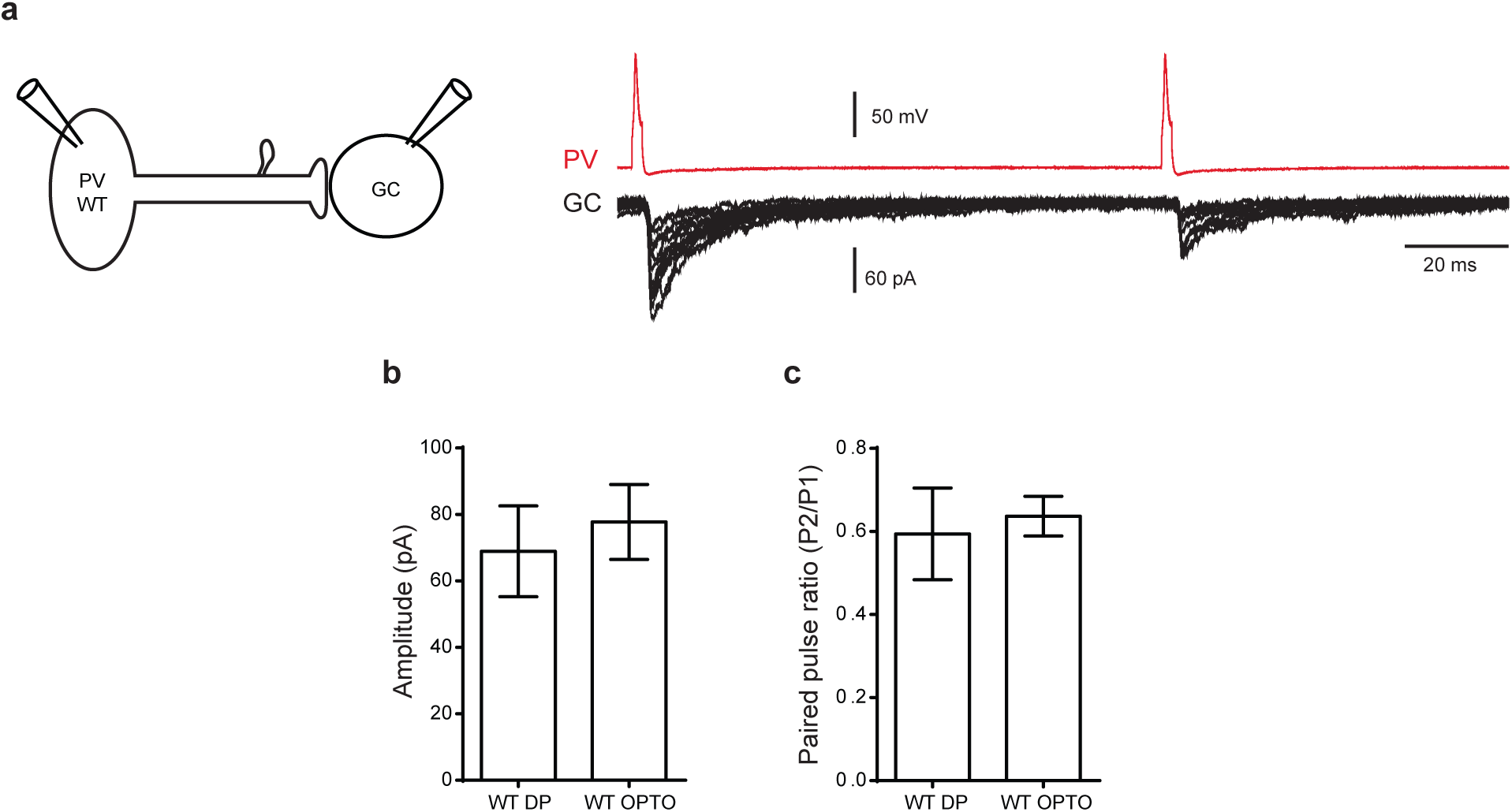
Comparison between double-patch and optogenetic monosynaptic responses in PV-GC synapses. **a.** Representative traces of paired-pulse (100 ms interval) double patch recordings from PV and GC neurons in WT mice. The monosynaptic GABAergic responses were recorded in GCs upon action potential generation in PV neurons (1-2 nA, 1ms) and in the presence of 50 µM APV and 10 µM CNQX to block glutamatergic receptors. **b, c**. Histograms showing that neither the 1^st^ amplitude (**b**) nor the paired-pulse ratio (**c**) of the GC responses evoked in the double patch configuration are different compared to the GC responses evoked by the optogenetic stimulation of PV neurons (n = 2 pairs for double patch recordings. Optogenetic data are shown also in Figure 4). Unpaired *t*-tests.

**Supplementary Figure S7.**
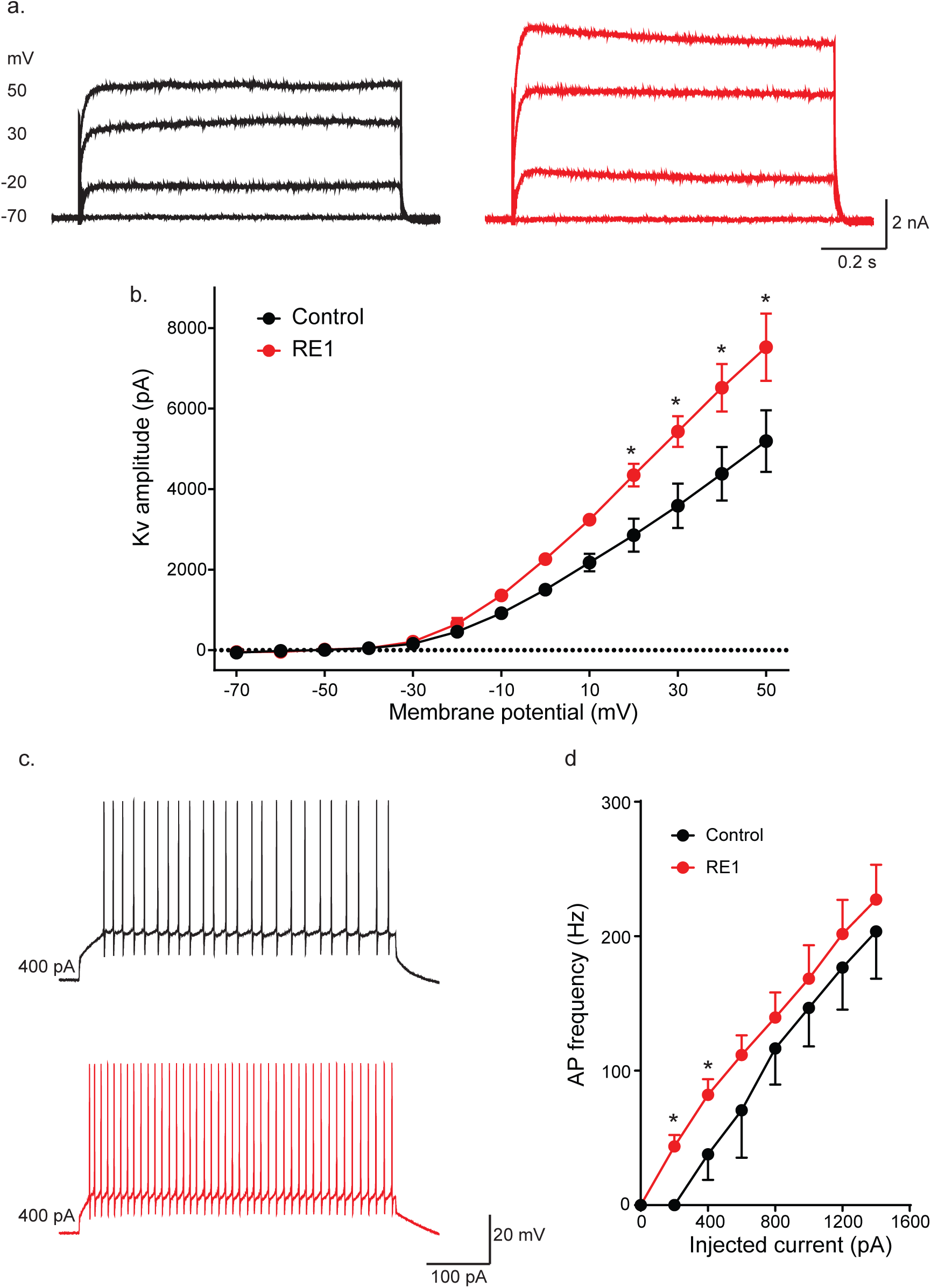
RE1 increases the amplitude of Kv currents and the firing of Purkinje neurons. **a.** Representative traces of Kv potassium currents evoked with 10 mV potential steps from −70 to +50 mV in cerebellar Purkinje neurons in control (black) and after the addition of 1µM RE1 (red). **b.** RE1 (1µM) significantly increased the amplitude of the Kv current in cerebellar Purkinje neurons (n = 3 neurons; paired *t*-test, \**p* < 0.05). **c.** Representative traces of action potential (AP) firing in cerebellar Purkinje neurons in response to 400 pA injected current in control (black) and after the addition of 1µM RE (red). **d.** RE1 (1µM) significantly increased the frequency of APs in cerebellar Purkinje neurons (n = 3 neurons; paired *t*-test, \**p* < 0.05).

## Materials and Methods

For all experiments, authenticated reagents were commercially purchased from local vendors.

### Synthesis of RE1

RE1, **1**, was prepared starting with the commercially available starting materials, as described in the patent literature^1^ with minor modification (Scheme S-1). All commercial chemicals and solvents were reagent grade and used without further purification. Air-sensitive reactions were performed under argon protection. Analytical thin layer chromatography was performed on Merck 250 μM silica gel F254 plates, and preparative thin layer chromatography on Merck 1000 μM silica gel F254 plates obtained from EMD Millipore corporation. Column chromatography was performed using CombiFlash instrument and Silica gel SNAP columns. The identity of each product was determined using NMR (Bruker 600 MHz instrument) and Agilent LC-MS. Chemical shifts are reported in δ values in ppm downfield from TMS as the internal standard. ^1^H data are reported as follows: chemical shift, multiplicity (s = singlet, d = doublet, t = triplet, q = quartet, m = multiplet), coupling constant (Hz), integration. Purity of final compound was determined using Agilent 1260 Infinity Series II preparative high-pressure liquid chromatography (HPLC) equipped with 1260 Infinity II multiple wavelength detector and Gemini® 5 µm C18 110 Å LC column (100 × 4.6 mm), mobile phase: Acetonitrile and water (0.1% trifluoroacetic acid), 80:20 isocratic. Compound **1** (RE1) was found more than 95% pure, as analyzed by LC-MS (Figure S-1).

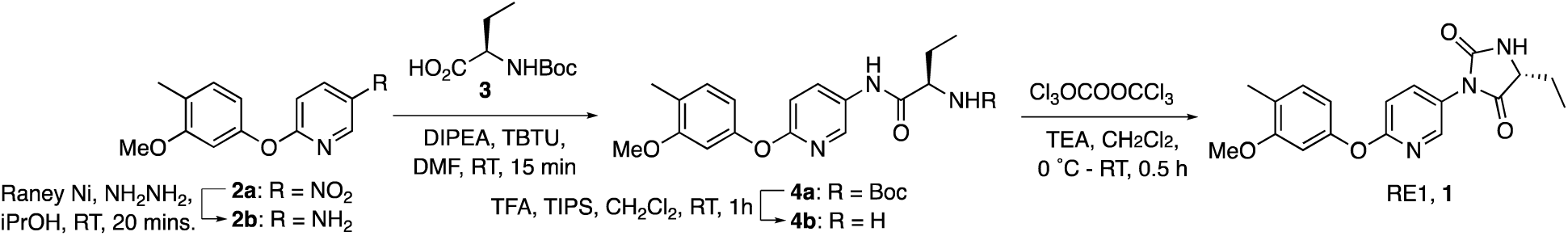

**Figure S-1.**
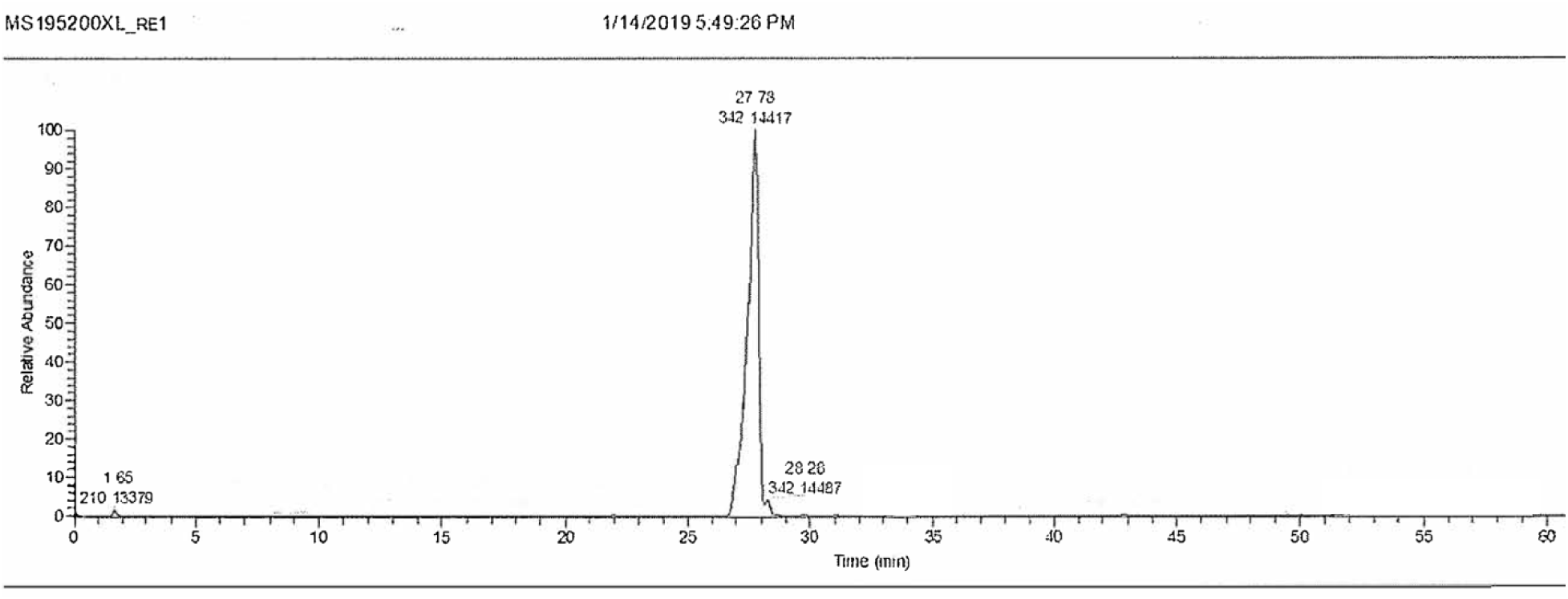
LC trace of RE1. Compound 1 was analyzed using LC-MS showing a major peak (>95%) for compound **1** (RE1). LC of compound **1** was performed on Agilent instrument as described above in methods.

#### Scheme S-1

Synthesis of RE1.

#### Compound 2a

A mixture of 4-methyl-3-methoxyphenol (1.97 g, 14.25 mM), 2-chloro-5-nitropyridine (2.71 g, 17.1 mmol) and K2CO3 (5.90 g, 42.8 mmol) in dry DMF (50 mL) was stirred at 115 °C for 2 hours. The reaction was worked-up using EtOAc and water. The organic layer was washed with brine, dried over anhyd. Na2SO4, and filtered. The organic layers were concentrated under vacuum and the residue was purified by silica gel chromatography using Hexanes-EtOAc (100:0 → 80:20) to afford compound **2a** (1.59 g, 43% Yield) as a light-yellow oil along with some impure fraction (930 mg, 25%). ^1^H-NMR (600 MHz, DMSO-d6): δ 9.02 (d, *J* = 2.8 Hz, 1H), 8.58 (dd, *J* = 9.1, 2.9 Hz, 1H), 7.18 (t, *J* = 7.6 Hz, 2H), 6.82 (d, *J* = 2.1 Hz, 1H), 6.68 (dd, *J* = 8.0, 2.2 Hz, 1H), 3.74 (s, 3H), 2.14 (s, 3H); MS (ESI): *m/z* 261.0866 [M+H]^+^

#### Compound 2b

To a solution of **2a** (710 mg, 2.73 mmol) in *iso*-propanol and hydrazine (9:1, 27 mL) was added Raney-Ni slurry (50% w/w in H2O, 710 µL), and the mixture was stirred at RT for 20 minutes. After the conversion was complete, as assessed by TLC, clear solution was obtained by filtration and concentrated under the reduced pressure, and the residue was chromatographed over Silica gel and Hexanes-EtOAc (100:0 → 50:50) to afford **2b (**490 mg, 78% Yield) as a light-yellow oil. ^1^H-NMR (600 MHz, DMSO-d6): δ 7.53 (d, *J* = 2.8 Hz, 1H), 7.05 (dd, *J* = 8.8, 2.9 Hz, 1H), 7.03 (d, *J* = 7.4 Hz, 1H), 6.70 (d, *J* = 8.6 Hz, 1H), 6.57 (d, *J* = 2.1 Hz, 1H), 6.36 (dd, *J* = 8.0, 2.2 Hz, 1H), 5.05 (s, 1H), 3.71 (s, 3H), 2.08 (s, 3H); MS (ESI): *m/z* 231.1134 [M+H]^+^.

#### Compound 4a

To a solution of **2** (127 mg, 0.63 mmol) in dry DMF (2 mL) were added DIPEA (182 µL, 1.25 mmol) followed by TBTU (216 mg, 0.82 mmol). After the reaction mixture was stirred for 15 minutes at room temperature, intermediate **2b** (120 mg, 0.52 mmol) was added and stirring was continued overnight at the same temperature. The reaction mixture was quenched using ice. The resulting crude product was dissolved in CH2Cl2, dried over Na2SO4, filtered and solvents were removed under the reduced pressure. The residue was purified by chromatography over Silica gel using hexanes-ethyl acetate (1:0 → 3:2) to afford compound **4a** (210 mg, 97% Yield) as a white solid. ^1^H NMR (600 MHz, DMSO-d6): δ 10.10 (s, 1H), 8.32 (s, 1H), 8.05 (dd, *J* = 8.8, 2.6 Hz, 1H), 7.11 (d, *J* = 8.0 Hz, 1H), 7.01 (d, *J* = 7.6 Hz, 1H), 6.95 (d, *J* = 8.8 Hz, 1H), 6.69 (s, 1H), 6.53 (d, *J* = 7.8 Hz, 1H), 3.97 (q, *J* = 5.9 Hz, 1H), 3.73 (s, 3H), 2.12 (s, 3H), 1.68 (m, 1H), 1.60 (m, 1H), 1.38 (s, 9H), 0.89 (t, *J* = 7.2 Hz, 3H); MS (ESI): *m/z* 416.2708 [M+H]^+^.

#### Compound 4b

TFA (1.25 mL) was added to ice-cold solution of **4a** (210 mg, 0.50 mmol) in dry CH2Cl2 (5 mL) followed by TIPS (125 µl), and the reaction mixture was stirred at room temperature for 1 hour. Solvents and excess reagents were removed under the reduced pressure and the residue was purified by chromatograph over Silica gel column using CH2Cl2-MeOH (1:0 → 4:1) to afford **3b** (175 mg) as a colorless solid. Intermediate **4b** was taken to next step after MS analysis. MS (ESI): *m/z* 316.2424 [M+H]^+^.

#### Compound 1, RE1

A solution of triphosgene (74 mg, 0.25 mmol) in 3 mL dry CH2Cl2 was added slowly to a solution of intermediate **4b** (171 mg, 0.54 mmol) and TEA (394 µL, 2.83 mmol) in dry CH2Cl2 (10 mL) at ice-water temperature. After the reaction mixture was stirred for 30 minutes at the same temperature it was quenched with water (3 mL) and worked-up using CH2Cl2 and water. The combined organic layers were dried over Na2SO4 and concentrated under the reduced pressure. The residue was purified by PTLC using hexanes-EtOAc (1:1) as the mobile phase to afford the title compound **1** (RE1, 56 mg, 30% Yield) as a white solid. ^1^H NMR (600 MHz, CDCl3): δ 8.24 (d, *J =* 2.46 Hz, 1H), 7.70 (dd, *J =* 8.8, 2.6 Hz, 1H), 7.12 (d, *J =* 7.7 Hz, 1H), 6.95 (d, *J =* 8.8 Hz, 1H), 6.63 (dd, *J =* 9.5, 2.0 Hz, 1H), 6.62 (s, 1H), 5.44 (s, 1H), 4.19 (t, *J =* 5.5 Hz, 1H), 3.78 (s, 3H), 2.19 (s, 3H), 1.98 (m, 1H), 1.92 (m, 1H), 1.05 (t, *J =* 7.4 Hz, 3H). MS (ESI): *m/z* 342.1442 [M+H]^+^.

### Cell culture

For cell transfection studies, 80% confluent N2A cells (1 × 10^5^ cells / well) were transfected with 3 μg of either mouse Kv3.1β or HA-tagged Kv3.1α plasmid, and 3 μg of a plasmid expressing rat p11.

### Kv3.1β-GFP stably transfected cells and colocalization imaging

GFP-tagged mouse Kv3.1β (C-term) was inserted into pEZ-M68 plasmid (Gencopoeia), and was transfected to N2A cells growing on OptiMEM medium with puromycin (1.25 µg/ ml). Individual clones were identified by AX10 fluorescent microscope (Zeiss) and were manually isolated using cloning cylinders. In order to inspect individual cells, the selected clone was mixed with untransfected N2A cells and the mixed culture was grown on complete medium without the selection marker. For siRNA studies, 70% confluent N2A cells in 60 mm dish were transfected with 200 pmol of either p11 silencing siRNA oligos with the double stranded RNA sequences: CAU GGA ACG GGA GUU CCC UGG GUU U, AAA CCC AGG GAA CUC CCG UUC CAU G, (ThemoFisher Scientific, 10620319/ 293344 A03), or scramble oligos as negative control 24 hours after plating. To determine the subcellular localization of Kv3.1β-GFP, 1.0-2.0 × 10^5^ cells were plated in wells with BioCoat 12mm coverslips (Corning), and were transfected with either p11 siRNA constructs or scramble oligos. For labelling the golgi or the ER, cells were fixed with 4% PFA after 24 hours, whereas methanol was used for membrane labeling. Antibodies included rabbit anti GM 130 (Abcam 52649, 1:1000), rabbit anti calnexin (Abcam 22595, 1:1000), rabbit anti sodium potassium ATPase (Abcam 76020, 1:1000) and mouse anti GFP (Abcam 1218, 1:500). Detection was performed using a confocal LSM710 microscope (Zeiss) and colocalization coefficients were determined using Zen 2012 SP1 software (Zeiss).

### Animals

All procedures were approved by the Animal Care and Use Committee of the Rockefeller University. Animals were maintained C57/Bl6N mice and were housed in a 12-hour light/ dark interval with food and water *ad-libitum*. p11 cKO mice were generated by crossing p11 floxed^2–5^ with Pvalb^tm1^(cre)^Arbr/J^ mice and their Cre^+^ offspring were used for all studies with Cre^−^ littermates as WT control. PV*^TRAP^* and p11 cKO*^TRAP^* mouse lines were generated by crossing these mice with mice expressing loxP-stop-loxP-EGFP-RPL10a sequence in the Eef1α1 promoter (EEF1A1–LSL.EGFPL10)^6^. TRAP qPCR analysis including mRNA isolation, cDNA amplification and qPCR analysis of mRNA level were previously described ^5^, by pooling 8 hippocampi from 4 mice per sample.

### AAV preparation and stereotaxic delivery

The sequence of the mouse Kv3.1β gene was inserted into pAAV.Flex plasmid and the sequence was validated by sequencing. rAAV2/5.Flex.Kv3.1β, rAAV2/5.Flex.GFP, rAAV2/5DIO.mCherry, rAAV2/5DIO.Gi-DREADD, rAAV2/5DIO.Gis-DREADD, AAV9DIOYFP/ChETA, were packaged at or purchased from the Virus Vector Core Facility, UNC (Chapel Hill, NC). 8-12 weeks old mice were injected with 1μl of AAV to the ventral DG 3 weeks before the behavioral tests or physiological recordings. Coordinates were ±2.00, −2.92 and −2.20 mm lateral, posterior and ventral relative to Bregma, according to the Franklin and Paxinos Mouse Brain atlas, 3rd edition.

### Western Blot

Cells in culture were lysed in RIPA buffer 48 hrs post transfection. The hippocampus was freshly harvested and lysed in 2% SDS. Protein concentration was determined using BCA (Thermo Fisher Scientific, Waltham, MA). 20 μg protein was loaded onto 4-12% BisTris gel, and transferred onto PVDF membranes that were incubated in 5% Milk in TBST and then with mouse anti-Kv3.1β (NeuroMab UC-DAVIS, 75-041, 1: 1,000), rabbit anti-HA (Cell Signaling, 3724 1: 1,000) rabbit anti-β-actin (Cell Signaling Technologies, 4970, 1: 2,000) or goat anti-p11 (R&D Systems, AF2377,1:200).

### Electrophysiology

Mice between 8 and 12 weeks of age were euthanized with CO2. After decapitation and removal of the brains, transversal slices (400 μm thickness) were cut using a Vibratome 1000 Plus (Leica Microsystems, USA) at 2 °C in a NMDG-containing cutting solution (in mM): 105 NMDG (N-Methyl-D-glucamine), 105 HCl, 2.5 KCl, 1.2 NaH2PO4, 26 NaHCO3, 25 Glucose, 10 MgSO4, 0.5 CaCl2, 5 L-Ascorbic Acid, 3 Sodium Pyruvate, 2 Thiourea (pH was around 7.4, with osmolarity of 295–305 mOsm). After cutting, slices were left to recover for 15 minutes in the same cutting solution at 35 °C and for 1 h at room temperature (RT) in recording solution (see below). Whole-cell patch-clamp recordings were performed with a Multiclamp 700B/Digidata1550A system (Molecular Devices, Sunnyvale CA, USA). EGFP-positive PV neurons were selected for recording based on the expression of the fluorescent marker using an upright Olympus BX51WI microscope equipped with the appropriate filters (Olympus, Japan) and a SPECTRA X LED light engine (Lumencor, OR, USA). The extracellular solution used for recordings contained (in mM): 125 NaCl, 25 NaHCO3, 2.5 KCl, 1.25 NaH2PO4, 2 CaCl2, 1 MgCl2 and 25 glucose (bubbled with 95% O2 and 5% CO2). The slice was placed in a recording chamber (RC-27L, Warner Instruments, USA) and constantly perfused with oxygenated aCSF at 24 °C (TC-324B, Warner Instruments, USA) at a rate of 1.5–2.0 ml/min.

For measuring the Kv we used whole-cell voltage-clamp recordings from PV neurons. The intracellular solution contained (in mM): 126 K-gluconate, 4 NaCl, 1 MgSO4, 0.02 CaCl2, 0.1 BAPTA, 15 glucose, 5 HEPES, 3 ATP, 0.1 GTP (pH 7.3). Voltage steps of 10 mv (from −70 mV to 50 mV; 1s, every 10 s) were used to determine current-voltage (I-V) relationships. A P/4 protocol was used to eliminate leak currents. The amplitude of the current was measured on the steady-state part of the response. Tetradotoxin (TTX, 0.3 μM) was added in the bath to block Na+ currents.

For measuring the HCN currents we used whole-cell voltage-clamp recordings from PV neurons. The intracellular solution contained (in mM): 126 K-gluconate, 4 NaCl, 1 MgSO4, 0.02 CaCl2, 0.1 BAPTA, 15 glucose, 5 HEPES, 3 ATP, 0.1 GTP (pH 7.3). Voltage steps of 10 mv (from −120 mV to −70 mV; 1s, every 10 s) were used to determine current-voltage (I-V) relationships. A P/4 protocol was used to eliminate leak currents. The amplitude of the HCN current was measured as the difference between amplitude the initial current at the beginning of the voltage step and the amplitude of the current in the steady-state stage of the voltage step. Tetradotoxin (TTX, 0.3 μM) was added in the bath to block Na+ currents.

Miniature EPSCs were recorded in the presence of bicucculline and TTX in the K-gluconate internal solution. To measure miniature AMPA currents AP-5 was added to the drugs used above. Miniature IPSCs were recorded in the presence of CNQX, AP-5 and TTX in a Cl-rich internal solution (in mM): 126 KCl, 4 NaCl, 1 MgSO4, 0.02 CaCl2, 0.1 BAPTA, 15 Glucose, 5 HEPES, 3 ATP and 0.1 GTP in which the pH was adjusted to 7.3 with KOH and osmolarity was adjusted to 290 mosmol/l with sucrose.

For paired recordings the presynaptic PV neurons were patched with the intracellular K-gluconate solution while the postsynaptic granule cells were patched using the Cl-rich internal solution. The distance between neurons was between 50 and 100 µm. Once a stable whole-cell configuration was achieved for both cells, steps of current of 1-2 nA for 2-5 ms were injected in the presynaptic PV neurons to elicit an action potential. The respective GABAergic response in the granule cells was recorded in the presence of CNQX and AP-5. 50 responses were recorded for each pair of cells. Data were acquired at a sampling frequency of 100 kHz, filtered at 1 kHz and analyzed offline using pClamp10 software (Molecular Devices, Sunnyvale, CA, USA).

### Optogenetic recordings

Field light stimulation of ChETA-expressing PV neurons was done through a 40x objective using a SPECTRA X LED light engine (Lumencor, OR, USA). Evoked IPSCs were recorded in granule cells in the presence of CNQX and AP-5 (to block glutamatergic transmission) in a Cl-rich internal solution (in mM): 126 KCl, 4 NaCl, 1 MgSO4, 0.02 CaCl2, 0.1 BAPTA, 15 Glucose, 5 HEPES, 3 ATP and 0.1 GTP in which the pH was adjusted to 7.3 with KOH and osmolarity was adjusted to 290 mosmol/l with sucrose. Data were acquired at a sampling frequency of 100 kHz, filtered at 1 kHz and analyzed offline using pClamp10 software (Molecular Devices, Sunnyvale, CA, USA).

### Immunohistochemistry

Mice were transcardially perfused with 4°c cold 4% paraformaldehyde. Staining of 80 μm thick floating sections used commercial antibodies against Kv3.1β (NeuroMab UC-DAVIS, 75-041, 1: 500), PV (Sigma Aldrich, P0388, 1:2000) and GFP (Abcam 1218, 1:500), with secondary Alexa goat anti-mouse or goat anti-rabbit were used (Thermo Fisher Scientific, Waltham, MA). Auto fluorescence was used to detect mCherry and cell nuclei were detected using DRAQ5 (Thermo Fisher Scientific, Waltham, MA). 4-6 coronal sections of 80 μm thickness were stained per antibody per mouse. To quantify the number of PV cells co-expressing mCherry a total of 17.3 ± 3.79 PV+ cells of the sub-granular zone of the ventral dentate gyrus were used per mouse.

### Mouse Behavior

In all DREADD experiments, CNO (Sigma Aldrich, C0832, 4 mg/ kg, in 100 μl saline) was injected intraperitoneally 30 minutes before the test. In the SSDS experiment CNO was injected before the first defeat. In the novel object recognition test (NOR), CNO was injected before the training phase. For intraperitoneal injections of RE1, the drug was dissolved in 10% 2-hydroxypropyl-β-cyclodextrin 5% DMSO.

Chronic social defeat stress (CSDS) was applied as previously described ^7^. In the subthreshold SDS paradigm (SSDS), each intruder mouse was subjected to 3 sessions of 2-minute physical defeat. Between each session, the intruder spent 15 minutes in the divided side of the cage. At the end of the third session the intruders were returned to group housing in their home cage. The social interaction test was conducted the next day, using unfamiliar aggressors. Open field (OF), sucrose preference (SPT), novelty suppressed feeding (NSF), forced swim (FST) and tail suspension (TST) tests were conducted as previously described ^5, 8^. In the elevated plus maze (EPM), mice were placed in the center of a plus sign comprised of two open and two enclosed arms (dimensions of each arm: 35 cm long, 5 cm wide, 15 cm high), elevated 40 cm above the floor. The time spent in the open and enclosed arms was automatically detected for 5 minutes using Ethnovision 7.0 (Noldus, Wageningen, the Netherlands). In NOR, mice were placed in the center of an arena with two identical objects (training exploration phase), and again 24h later with or one familiar and one novel object (recognition phase). The time spent interacting with each object was automatically detected for 10 minutes using Ethnovision 7.0 (Noldus, Wageningen, the Netherlands).

### Statistical analysis

All data are expressed as means ± s.e.m. Statistical analysis was performed using GraphPad Prism7. In all experiments, *p* < 0.05 was considered significant.

## References

1. Petty, F. & Sherman, A.D. Plasma GABA levels in psychiatric illness. J Affect Disord 6, 131–138 (1984).

2. Romeo, B., Choucha, W., Fossati, P. & Rotge, J.Y. Meta-analysis of central and peripheral gamma-aminobutyric acid levels in patients with unipolar and bipolar depression. J Psychiatry Neurosci 42, 160228 (2017).

3. Knable, M.B., et al. Molecular abnormalities of the hippocampus in severe psychiatric illness: postmortem findings from the Stanley Neuropathology Consortium. Molecular psychiatry 9, 609–620, 544 (2004).

4. Sequeira, A., et al. Global brain gene expression analysis links glutamatergic and GABAergic alterations to suicide and major depression. PLoS One 4, e6585 (2009).

5. Martisova, E., et al. Long lasting effects of early-life stress on glutamatergic/GABAergic circuitry in the rat hippocampus. Neuropharmacology 62, 1944–1953 (2012).

6. Engin, E., Benham, R.S. & Rudolph, U. An Emerging Circuit Pharmacology of GABAA Receptors. Trends Pharmacol Sci 39, 710–732 (2018).

7. Klausberger, T. & Somogyi, P. Neuronal diversity and temporal dynamics: the unity of hippocampal circuit operations. Science 321, 53–57 (2008).

8. Hu, H., Gan, J. & Jonas, P. Interneurons. Fast-spiking, parvalbumin(+) GABAergic interneurons: from cellular design to microcircuit function. Science 345, 1255263 (2014).

9. Marin, O. Interneuron dysfunction in psychiatric disorders. Nat Rev Neurosci 13, 107–120 (2012).

10. Magloczky, Z. & Freund, T.F. Impaired and repaired inhibitory circuits in the epileptic human hippocampus. Trends in neurosciences 28, 334–340 (2005).

11. Svenningsson, P., et al. Alterations in 5-HT1B receptor function by p11 in depression-like states. Science 311, 77–80 (2006).

12. Egeland, M., Warner-Schmidt, J., Greengard, P. & Svenningsson, P. Neurogenic effects of fluoxetine are attenuated in p11 (S100A10) knockout mice. Biol Psychiatry 67, 1048–1056 (2010).

13. Warner-Schmidt, J.L., et al. A role for p11 in the antidepressant action of brain-derived neurotrophic factor. Biol Psychiatry 68, 528–535 (2010).

14. Oh, Y.S., et al. SMARCA3, a chromatin-remodeling factor, is required for p11-dependent antidepressant action. Cell 152, 831–843 (2013).

15. Milosevic, A., et al. Cell- and region-specific expression of depression-related protein p11 (S100a10) in the brain. J Comp Neurol 525, 955–975 (2017).

16. Medrihan, L., et al. Initiation of Behavioral Response to Antidepressants by Cholecystokinin Neurons of the Dentate Gyrus. Neuron 95, 564–576 e564 (2017).

17. Rudy, B. & McBain, C.J. Kv3 channels: voltage-gated K+ channels designed for high-frequency repetitive firing. Trends Neurosci 24, 517–526 (2001).

18. Atzori, M., et al. H2 histamine receptor-phosphorylation of Kv3.2 modulates interneuron fast spiking. Nature neuroscience 3, 791–798 (2000).

19. Kaczmarek, L.K. & Zhang, Y. Kv3 Channels: Enablers of Rapid Firing, Neurotransmitter Release, and Neuronal Endurance. Physiological reviews 97, 1431–1468 (2017).

20. Hoppa, M.B., Gouzer, G., Armbruster, M. & Ryan, T.A. Control and plasticity of the presynaptic action potential waveform at small CNS nerve terminals. Neuron 84, 778–789 (2014).

21. Goldberg, E.M., et al. Specific functions of synaptically localized potassium channels in synaptic transmission at the neocortical GABAergic fast-spiking cell synapse. The Journal of neuroscience: the official journal of the Society for Neuroscience 25, 5230–5235 (2005).

22. Girard, C., et al. p11, an annexin II subunit, an auxiliary protein associated with the background K+ channel, TASK-1. EMBO J 21, 4439–4448 (2002).

23. Okuse, K., et al. Annexin II light chain regulates sensory neuron-specific sodium channel expression. Nature 417, 653–656 (2002).

24. van de Graaf, S.F., et al. Functional expression of the epithelial Ca(2+) channels (TRPV5 and TRPV6) requires association of the S100A10-annexin 2 complex. EMBO J 22, 1478–1487 (2003).

25. Donier, E., Rugiero, F., Okuse, K. & Wood, J.N. Annexin II light chain p11 promotes functional expression of acid-sensing ion channel ASIC1a. J Biol Chem 280, 38666–38672 (2005).

26. Boddum, K., et al. Kv3.1/Kv3.2 channel positive modulators enable faster activating kinetics and increase firing frequency in fast-spiking GABAergic interneurons. Neuropharmacology 118, 102–112 (2017).

27. Anacker, C., et al. Hippocampal neurogenesis confers stress resilience by inhibiting the ventral dentate gyrus. Nature 559, 98–102 (2018).

28. Zou, D., et al. DREADD in parvalbumin interneurons of the dentate gyrus modulates anxiety, social interaction and memory extinction. Curr Mol Med 16, 91–102 (2016).

29. Cheng, J., Umschweif, G., Leung, J., Sagi, Y. & Greengard, P. HCN2 Channels in Cholinergic Interneurons of Nucleus Accumbens Shell Regulate Depressive Behaviors. Neuron 101, 662–672 e665 (2019).

30. Lee, K.W., et al. p11 regulates the surface localization of mGluR5. Mol Psychiatry 20, 1485 (2015).

31. Kilisch, M., Lytovchenko, O., Schwappach, B., Renigunta, V. & Daut, J. The role of protein-protein interactions in the intracellular traffic of the potassium channels TASK-1 and TASK-3. Pflugers Arch 467, 1105–1120 (2015).

32. Oh, S.J., et al. Hippocampal mossy cell involvement in behavioral and neurogenic responses to chronic antidepressant treatment. Mol Psychiatry (2019).

33. Eriksson, T.M., et al. Bidirectional regulation of emotional memory by 5-HT(1B) receptors involves hippocampal p11. Mol Psychiatry (2012).

34. Lopez, A.J., et al. Promoter-Specific Effects of DREADD Modulation on Hippocampal Synaptic Plasticity and Memory Formation. J Neurosci 36, 3588–3599 (2016).

35. Warner-Schmidt, J.L., et al. Cholinergic interneurons in the nucleus accumbens regulate depression-like behavior. Proc Natl Acad Sci U S A 109, 11360–11365 (2012).

36. Hanada, Y., et al. p11 in Cholinergic Interneurons of the Nucleus Accumbens Is Essential for Dopamine Responses to Rewarding Stimuli. eNeuro 5(2018).

37. Schmidt, E.F., et al. Identification of the cortical neurons that mediate antidepressant responses. Cell 149, 1152–1163 (2012).

38. Hu, H., Roth, F.C., Vandael, D. & Jonas, P. Complementary Tuning of Na(+) and K(+) Channel Gating Underlies Fast and Energy-Efficient Action Potentials in GABAergic Interneuron Axons. Neuron 98, 156–165 e156 (2018).

39. Muona, M., et al. A recurrent de novo mutation in KCNC1 causes progressive myoclonus epilepsy. Nature genetics 47, 39–46 (2015).

40. Yanagi, M., et al. Kv3.1-containing K(+) channels are reduced in untreated schizophrenia and normalized with antipsychotic drugs. Molecular psychiatry 19, 573–579 (2014).

41. Sauer, J.F., Struber, M. & Bartos, M. Impaired fast-spiking interneuron function in a genetic mouse model of depression. eLife 4(2015).

42. Shen, S., et al. Schizophrenia-related neural and behavioral phenotypes in transgenic mice expressing truncated Disc1. The Journal of neuroscience: the official journal of the Society for Neuroscience 28, 10893–10904 (2008).

43. Sagi, Y., et al. Emergence of 5-HT5A signaling in parvalbumin neurons mediates delayed antidepressant action. Mol Psychiatry (2019).

## References

1. Alvaro, G., et al. Preparation of hydantoin derivatives as Kv3 channel inhibitors. 279pp. (Autifony Therapeutics Limited, UK. 2012).

2. Warner-Schmidt, J.L., et al. Role of p11 in cellular and behavioral effects of 5-HT4 receptor stimulation. J Neurosci 29, 1937–1946 (2009).

3. Warner-Schmidt, J.L., et al. Cholinergic interneurons in the nucleus accumbens regulate depression-like behavior. Proc Natl Acad Sci U S A 109, 11360–11365 (2012).

4. Virk, M.S., et al. Opposing roles for serotonin in cholinergic neurons of the ventral and dorsal striatum. Proc Natl Acad Sci U S A 113, 734–739 (2016).

5. Medrihan, L., et al. Initiation of Behavioral Response to Antidepressants by Cholecystokinin Neurons of the Dentate Gyrus. Neuron 95, 564–576 e564 (2017).

6. Stanley, S., et al. Profiling of Glucose-Sensing Neurons Reveals that GHRH Neurons Are Activated by Hypoglycemia. Cell Metab 18, 596–607 (2013).

7. Cheng, J., Umschweif, G., Leung, J., Sagi, Y. & Greengard, P. HCN2 Channels in Cholinergic Interneurons of Nucleus Accumbens Shell Regulate Depressive Behaviors. Neuron 101, 662–672 e665 (2019).

8. Sagi, Y., et al. Emergence of 5-HT5A signaling in parvalbumin neurons mediates delayed antidepressant action. Mol Psychiatry (2019).

